# PARP1 directly disassembles nucleosomes to regulate DNA repair

**DOI:** 10.64898/2026.03.22.713488

**Authors:** Ashish Verma, Changlei Zhu, Bernadette Truong, Silvija Bilokapic, Rebekah DeVries, Aaron Pitre, Anang A. Shelat, Mario Halic, Hai T. Dao

## Abstract

Upon DNA damage, chromatin remodeling is rapidly initiated to promote chromatin accessibility, thereby facilitating the recruitment and assembly of repair factors. Although this enhanced accessibility has been linked to poly(ADP-ribose) polymerase (PARP) activity, the mechanism by which cells overcome the nucleosome barrier remains unclear. Using our designer chromatin system, we uncovered a previously uncharacterized activity of PARP1, whereby it directly and asymmetrically evicts histone dimers proximal to DNA strand breaks from nucleosomes to generate oriented hexasomes. In the presence of HPF1, PARP1 generates stable PARylated hexasomes, an open chromatin intermediate that can serve as a bifunctional hub for recruitment of DNA- and PAR-dependent factors. Using cellular assays, we demonstrated that PARP activity is both required and sufficient to drive chromatin accessibility and the recruitment of repair factors, with direct involvement of subnucleosomal species. Unexpectedly, we identified the C-terminal tail of histone H2A, a motif harboring recurrent cancer-associated mutations, as a critical determinant of efficient PARP1-mediated nucleosome disassembly. Deletion of the H2A tail sensitizes cells to DNA-damaging agents and PARP inhibitors, implicating a functional role of PARP1-mediated nucleosome disassembly in DNA repair. Together, our findings support a model in which PARP1 directly drives histone eviction, leading to the formation of subnucleosomes that facilitate efficient DNA repair.

## Main

The compaction of eukaryotic genomes into chromatin poses a physical barrier to DNA-templated processes. To overcome this, cells have evolved multiple mechanisms that regulate chromatin structure by modulating nucleosomes^1–4^, the repeating units of chromatin. During the DNA damage response (DDR), both ATP-dependent chromatin remodeling complexes and histone-modifying enzymes are essential for facilitating the exposure of DNA lesions, making them accessible to repair factors^5–9^. Protein poly(ADP-ribosyl)ation (PARylation) is one of the earliest events during multiple DDR pathways, including single-stranded DNA (ssDNA) and double-stranded DNA (dsDNA) breaks ^10–14^. This post-translational modification is catalyzed by poly(ADP-ribose) polymerases (PARPs), with PARP1 being the most prominent contributor, accounting for approximately 90% of PARylation induced by DNA damage^10^.

PARP1 is a multidomain enzyme with central roles in chromatin regulation, including transcription regulation^15,16^, in addition to its well-established functions in DNA repair^10^. In the context of DDR, PARP1 performs two primary functions: it acts as a DNA lesion sensor through multiple DNA-binding domains^17,18^, and as a catalytic enzyme that mediates PARylation of itself (automodification) or other proteins (*trans*-PARylation), including histone linker H1 proteins, mainly at glutamic acid residues^18,19^. In the presence of the auxiliary histone PARylation factor 1 (HPF1), PARP1 can efficiently modify core histones and switch its modification specificity from glutamic acid to serine^20,21^.

PARylation of histone proteins modulates chromatin structure both in *cis* and *trans* mechanisms^22^. Owing to the highly negatively charged phosphate backbone of poly(ADP-ribose) (PAR), PARylation of histones weakens histone-DNA interactions, thereby disrupting nucleosome packaging and internucleosomal contact. This *cis* effect was first demonstrated in early biochemical studies by Poirier and Mandel^23–25^ and later substantiated by work from Muir^26^, establishing the model that PARylation promotes chromatin decompaction and enhances accessibility to facilitate downstream repair^18^. Subsequent studies combining laser microirradiation with live-cell imaging have reinforced this model, showing that PARP activity drives localized chromatin expansion and increased accessibility for transcription factors and chromatin-associated proteins at DNA damage sites^11,12,27–30^. Concomitant with chromatin expansion, linker histone H1 is rapidly depleted from DNA lesions, presumably due to electrostatic repulsion between negatively charged PAR chains on PARylated H1 and surrounding chromatin^28^. Notably, displacement of core histone H2A and H2B has also been observed, suggesting that PARP-dependent pathways can overcome the nucleosome barrier by evicting the core histones to contribute to chromatin accessibility^28,31,32^.

In addition to the *cis*-effect, PARylated chromatin also functions as a recruiting hub to elicit the *trans*-effect on chromatin, as an expanding repertoire of PAR-reading proteins has been identified and shown to localize to the DNA lesion^33–37^. Among these are the ATP-dependent chromatin remodeler ALC1, which has been shown to facilitate nucleosome repositioning away from damage sites^38–41^. However, ALC1 has not been shown to mediate nucleosome eviction or disassembly. Although the BAF complex^42–46^ and INO80 complex^47–52^, which possess nucleosome disassembly activity, have been implicated in the response to dsDNA breaks, their recruitment kinetics do not align with the rapid PARP1-dependent histone displacement observed immediately following DNA damage^28^. Together, there is a key gap in our understanding of how nucleosomes are disassembled during the early stage of DNA repair.

The nucleosome is assembled from a (H3–H4)₂ tetramer and two H2A–H2B dimers, which together form a C2-symmetric histone octamer around which ∼147 bp of DNA is wrapped^53^. The removal of one or two histone H2A-H2B dimers from the nucleosome results in the formation of hexasomes and tetrasomes, respectively^54–59^. These subnucleosomal species have been implicated in DNA repair^60^, in addition to their roles in transcription and DNA replication^54,58,61–70^. Even though PARP-dependent histone displacement occurs rapidly following DNA damage^28,31,32^, it remains unclear whether PARP enzymes directly drive nucleosome disassembly or instead act indirectly through recruitment of downstream ATP-dependent chromatin remodeling complexes^5,9,30,71,72^.

In this study, we combined biochemical and cellular assays to dissect the functions of PARP1 in chromatin remodeling. Our results revealed a previously uncharacterized NAD^+^-dependent chromatin-remodeling mechanism in which PARP1 asymmetrically disassembles nucleosomes to generate oriented hexasomes. By probing changes in cellular chromatin structure at the early stage of DNA repair, we demonstrated that PARP1 is both necessary and sufficient to promote chromatin accessibility, with direct involvement of subnucleosomal intermediates. Further studies revealed that the H2A C-terminal tail serves as a regulatory motif for PARP1-mediated chromatin disassembly and is required for cell growth under DNA-damage conditions. Notably, several highly recurrent oncohistones were observed at this tail, underscoring the relevance of PARP1-mediated nucleosome disassembly to disease-associated chromatin dysfunction^73^. Collectively, our studies suggest a model in which PARP1 activity directly promotes nucleosome disassembly, thereby enhancing chromatin accessibility at DNA lesions and promoting efficient DNA repair.

## Results

### PARP1 directly and asymmetrically disassembles nucleosomes to generate the oriented hexasomes in an NAD^+^ dependent manner

Automodified PARP1 has been reported to destabilize histone-DNA interactions through its negatively charged poly(ADP-ribose) (PAR) chains^74^. We therefore hypothesized that autoPARylated PARP1 might similarly promote nucleosome disassembly. Because DNA strand breaks occur randomly throughout the genome, lesions frequently arise adjacent to nucleosomes and can be recognized by PARP1 (Fig 1A, left). To model this configuration, we strategically designed mononucleosome substrates that allow us to monitor PARP1-dependent activity on one side while restricting activity on the other (Fig 1A, right).

**Figure 1.**
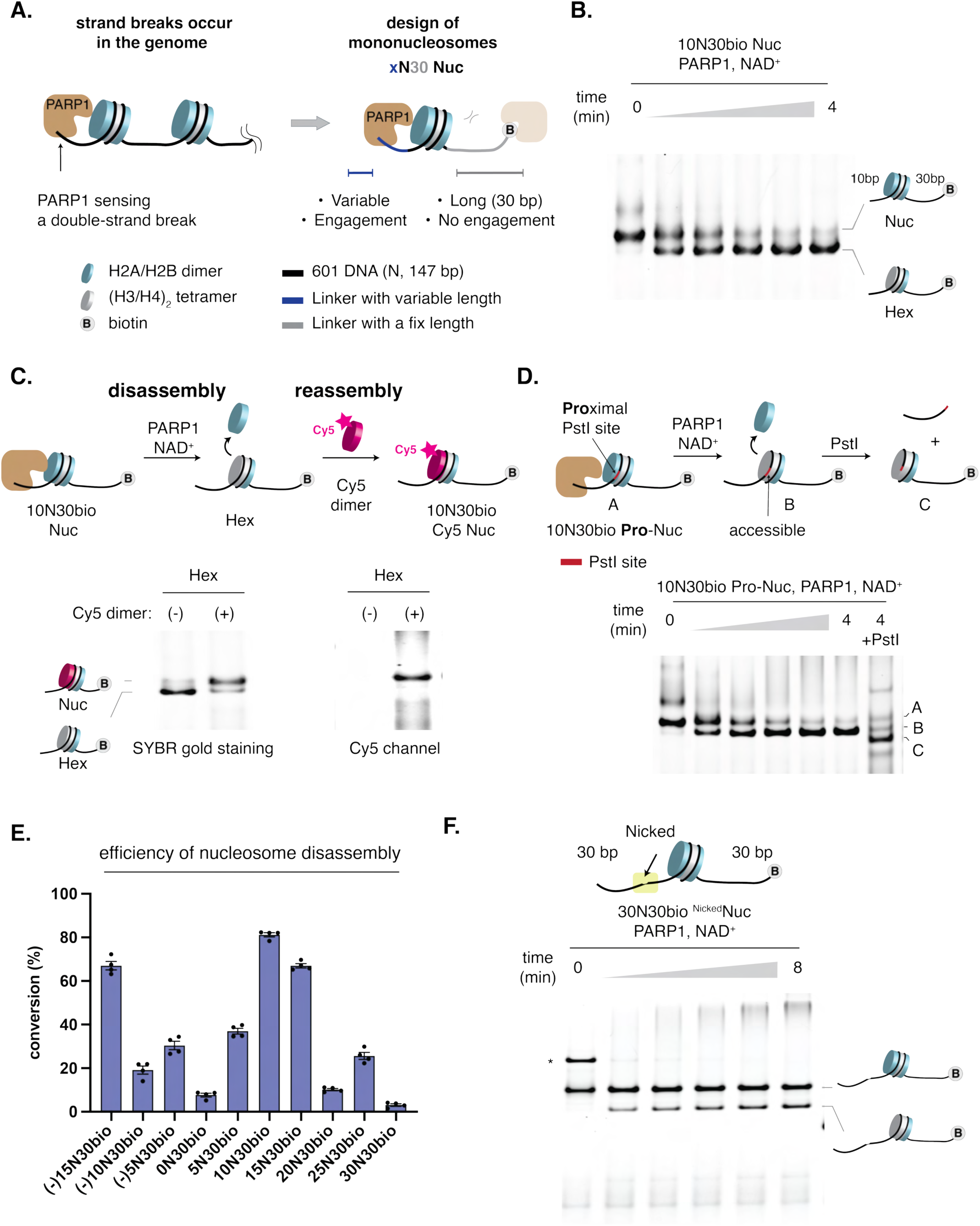
PARP1 asymmetrically disassembles nucleosomes to generate oriented hexasome intermediates in biochemical assays. (A) Design of mononucleosome substrates to model PARP1 activity on chromatin bearing DNA strand breaks. (Left) Model illustrating PARP1 binding to a DNA double-strand break and interacting with the adjacent nucleosome. (Right) Diagram of chromatin substrates containing a fixed 30-bp biotinylated linker and a variable linker. The long (30 bp) linker restricts nucleosome engagement from that side, allowing assessment of linker-length–dependent PARP1-mediated chromatin remodeling. (B) NAD^+^-dependent PARP1 activity facilitates nucleosome disassembly, as monitored in an electrophoretic mobility shift assay (EMSA). NAD⁺ was added to initiate the reaction. At the indicated time points, aliquots were taken and quenched with 60 bp DNA fragments to sequester PARP1 from chromatin, enabling tracking of chromatin structural states during the reaction. Gel was stained with SYBR™ Gold to visualize total DNA. (C) Nucleosome reassembly assay demonstrating incorporation of a Cy5-labeled histone dimer into hexasome intermediates. After quenching the disassembly reaction (Fig 1B, t = 4 min) with dsDNA fragments and olaparib (12.5 μM), Cy5-labeled histone dimers were added and incubated at 30 °C to allow nucleosome reassembly. The gel was imaged in the Cy5 channel to detect Cy5 incorporation, followed by SYBR™ Gold staining to visualize total DNA. (D) The histone dimer proximal to the 10 bp linker is selectively evicted, exposing the PstI enzyme site to become accessible to digestion by PstI. Gel was stained with SYBR™ Gold to visualize total DNA. (E) Linker length-dependent nucleosome disassembly quantified at t = 2 minutes. (F) PARP1-mediated nucleosome disassembly triggered by a nick positioned 10 bp from the nucleosome entry site. Gel was stained with SYBR™ Gold to visualize total DNA. N, nucleosome assembled on 147-bp Widom 601 DNA; NAD^+^, nicotinamide adenine dinucleotide. Nucleosome disassembly reaction conditions: nucleosome (0.1 μM), PARP1 (0.2 μM), NAD+ (0.5 mM).

PARP1 is known to bind broken DNA ends while engaging with chromatin; therefore, the linker length, representing the distance between the nucleosome core and the DNA break, is likely to influence its activity on nucleosomes^17,19^. To systematically investigate PARP1-mediated nucleosome disassembly, we generated a series of designer nucleosomes using Widom 601 DNA (i.e., N) fragments containing a long, fixed 5’-biotinylated DNA overhang on one side (30 bp) and a variable-length overhang on the other (x bp), hereafter referred to as xH30bio nucleosomes (Fig. 1A, right). The long 30 bp overhang restricts PARP1 interaction with the nucleosome when binding to the DNA end on that side, enabling us to assess the impact of PARP1 on chromatin as a function of the variable overhangs on the variable side. To further mitigate the effects of the longer linker, we included a biotin group at the 5’ end and installed the streptavidin blockage when applicable (Fig. S1A). Analogously, we assembled synthetic hexasome substrates containing a 601-derived high-positioning sequence (H), hereafter referred to as xH30bio hexasomes (Fig. S1B)^54^.

To track chromatin status over time, we utilized dsDNA fragments to sequester PARP1 from chromatin in an electrophoretic mobility shift assay (EMSA) (Fig. S1C, lanes 1 and 2). In our model 10N30bio nucleosome, upon treatment of NAD^+^, we observed the formation of stable, fast-moving chromatin species (lower bands, Figs. 1B, S1C), concomitant with automodification of PARP1, without any significant modification of histones (Fig. S1D). Native gel electrophoresis followed by western blot analysis revealed the presence of both H3 and H2B on the fast-moving species, suggesting a hexasomal structure (Figs. S1E, F). Furthermore, the mobility shift of this chromatin species matched with the standard oriented hexasome prepared using a previously described method (Fig. S1G)^54,58,59^. Consistently, in a nucleosome reassembly assay in which a Cy5-labeled histone dimer was added to reactions containing the putative hexasome species, we observed reassembly of labeled nucleosomes (Fig. 1C)^58,59^. Nucleosome disassembly was also observed when using streptavidin blockage at the fixed 30bp linker, suggesting that PARP1 engages the variable linker side, with the length of 10bp in this case, to affect chromatin remodeling (Fig. S1H). The enhanced reaction rate with the streptavidin-blocked nucleosome likely results from increased effective PARP1 concentration at the flexible DNA end when the fixed side is blocked (Figs. S1C vs. S1H).

The nucleosome has two dimers that can undergo disassembly during NAD^+^-dependent PARP1-mediated remodeling. To test the specificity of nucleosome disassembly, we prepared two 10N30bio nucleosome substrates: a PstI-proximal nucleosome (Pro-Nuc) and a PstI-distal nucleosome (Dis-Nuc). Each substrate contained a PstI recognition site embedded within nucleosomal DNA contacting the histone dimer located either proximal or distal to the 10 bp linker (Figs. 1D and S1I). Using EMSA followed by PstI restriction enzyme digestion, digested hexasomes were detected exclusively in 10N30bio Pro-Nuc, supporting asymmetric nucleosome disassembly (Figs. 1D and S1I). To control for the asymmetry of the 601 DNA sequence, we prepared and tested bio30N10 nucleosome substrates (bio30N10 Pro-Nuc and bio30N10 Dis-Nuc) and observed a similar trend (Figs. S1J,K)^58^. Consistent with previous studies on histone-DNA complexes, we observed the regeneration of nucleosomal species upon treatment with poly(ADP-ribose) glycohydrolase (PARG) (Fig. S1L, lanes 4 vs 5) ^25,74^.

Systematic studies revealed a linker length-dependent trend, with the maximum activity observed when the linkers are approximately 10-15 bp in length (Figs. 1E, S1M-O). While long overhangs (e.g., 30 bp) are expected to disfavor the PARP1-chromatin engagement, short overhangs (e.g., 0 bp) might result in a steric clash between the nucleosome core and PARP1, hence negatively impacting the activity. Interestingly, the remodeling activity increased as the flexible linker side was truncated into nucleosomes, corresponding to the double-strand break that occurs within the nucleosomes (Figs. 1E, S1P,Q). Notably, the nicked nucleosome also disassembled under similar conditions, suggesting that single-stranded breaks can also trigger PARP1-mediated chromatin remodeling (Figs. 1F vs. S1R). Intriguingly, we did not observe nucleosome disassembly with PARP2, even with substrates carrying a 5’ phosphate, suggesting that the reaction is more specific to PARP1 (Figs. S1S and T). While nucleosomes undergo disassembly, the hexasome counterparts with a damaged side proximal to the loss dimer remain stable under similar conditions (Figs. S1U-W). Importantly, when we switched the 10 bp-linker to the distal side relative to the loss dimer, we observed hexasome disassembly, suggesting that nucleosome remodeling is more specific to the removal of histone dimers (Figs. S1X,Y).

The 601 sequence is an engineered high-affinity nucleosome-positioning DNA sequence that can influence disassembly. To control for this synthetic effect, we assembled nucleosomes using native DNA sequences containing the 5S rDNA or Lin28B genes^75^. Our data indicate that nucleosome disassembly occurs more efficiently on native DNA substrates, suggesting the generality of this activity in a cellular context (Fig. S1Z, S1AA, S1AB). Moreover, preactivated PARP1 does not disrupt nucleosomes, suggesting that nucleosome disassembly is coupled to direct PARP1-nucleosome engagement (Fig. S1AC). Collectively, our results revealed a previously uncharacterized NAD^+^-dependent chromatin-remodeling activity of PARP1, triggered by DNA lesions, that drives the generation of oriented hexasome intermediates.

### PARP1-mediated nucleosome disassembly facilitates the binding of repair factors to synthetic chromatin

H1 proteins are key chromatin components that interact with the majority of nucleosomes in most eukaryotic cells to form chromatosomes^76^. Given that PARP1 and H1 compete for nucleosome binding^77^ and display opposite occupancy patterns at regulatory regions^78^, we investigated whether PARP1 could promote core nucleosome disassembly in the presence of H1. To test this, we reconstituted a chromatosome complex using the 10N30bio nucleosome and histone H1.0, then subjected it to a PARP1-mediated nucleosome disassembly assay (Figs. 2A and S2A). Although the chromatosome band initially remained unchanged, we later detected the formation of a lower band corresponding to the hexasome intermediate (Fig. 2A). Increasing the concentration of H1.0 extended the duration of the initial lag phase, supporting a sequential mechanism where histone H1.0 is evicted before the disassembly of the core nucleosome (Figs. 2B and S2B). In line with this model, polyADP-ribosylated H1.0 was detected at an early time point (t = 4 minutes) prior to the nucleosome disassembly event (Figs. 2B and 2C). Additionally, the overlap in the timing of automodification of PARP1 and hexasome formation supports a mechanism in which PARylated PARP1 plays a role in the displacement of the histone H2A/H2B dimer (Figs. 2B and 2C). Notably, the remodeled chromatin treated with PARG reassembled into a chromatosome structure containing monoADP-ribosylated H1.0, consistent with the reversibility of PARP1-mediated chromatin structure alteration (Figs. 2A, 2B, and S2C). Hexasome formation was also observed with the chromatosome formed using H1.4, suggesting a general mechanism of chromatosome disassembly (Fig. S2D).

**Figure 2.**
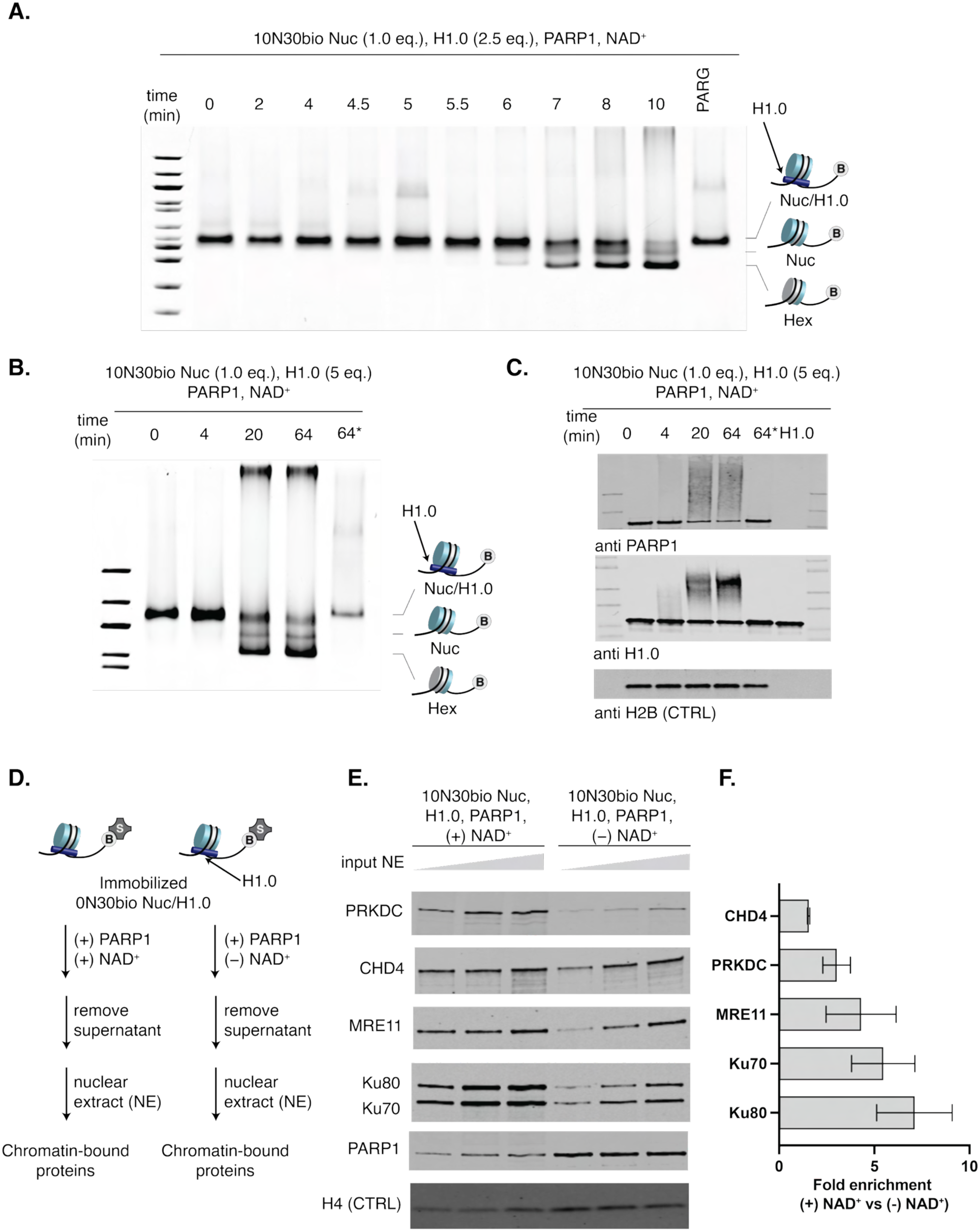
PARP1 disassembles chromatosomes to enhance chromatin accessibility. (A) Time-course experiment illustrating NAD⁺-dependent PARP1-mediated disassembly of nucleosome-H1.0 chromatosome complexes (lanes 2–11); 2.5 equivalents of H1.0 were used. Disassembled chromatin can be converted back to chromatosomes upon treatment with Poly (ADP-ribose) glycohydrolase (PARG) (lane 12). Gel was stained with SYBR™ Gold to visualize total DNA. (B, C) Time-course experiment illustrating NAD⁺-dependent PARP1-mediated chromatosome disassembly as monitored by native gel (B) and western blot (C); 5.0 equivalents of H1.0 were used. PARP1 and H1 undergo *trans*-PARylation to generate slower-migrating species, whereas core histone H2B remains unchanged. (D) Schematic of the chromatin pulldown assay with HeLa nuclear extract used to assess the effects of PARP1-driven chromatin disassembly. (E) PARP1-induced chromatin disassembly promotes recruitment of DNA repair factors, including Ku80, Ku70, PRKDC, MRE11, and CHD4. (F) Quantification of fold enrichment of repair factors in the presence of NAD⁺ relative to reactions lacking NAD⁺ in the experiment shown in Fig. 2E, under the lowest nuclear extract (NE) input conditions. Error bars represent SEM from two independent replicates. Chromatin disassembly reaction conditions: nucleosome (0.1 μM), H1.0 (0.25 or 0.5 μM), PARP1 (0.2 μM), NAD+ (0.5 mM).

The disassembly of nucleosomes and chromatosomes results in the release of approximately 35-50 base pairs of unbound DNA, potentially providing binding sites for DNA repair factors. To investigate the functional impact of PARP1-mediated chromatin remodeling, we performed a chromatin affinity pulldown assay using HeLa S3 nuclear extract with chromatosome and PARP1-induced remodeled chromatosome baits (Figs. 2D-F, S2E-G). Our results showed that both components of the Ku70/80 heterodimer, the key initiator of the non-homologous end joining (NHEJ) pathway, and their associated factor PRKDC were enriched on remodeled chromatin. Similarly, MRE11, a core component of the MRN complex (MRE11, RAD50, NBS1) involved in homologous recombination (HR), was also enriched. Consistent with previous studies, CHD4, a repair factor with DNA-binding capabilities, showed enhanced binding to remodeled substrates^27^. We also found that the trend is less pronounced in the absence of H1 proteins, suggesting that chromatosomes are less accessible to DNA repair factors (Figs. S2H-J). Altogether, these findings suggest that PARP1-mediated chromatin disassembly promotes a more accessible chromatin structure, thereby facilitating the recruitment of DNA repair factors.

### PARP1/HPF1 generates PARylated hexasomes in biochemical assays

Since histone PARylation factor 1 (HPF1) acts as a PARP1 cofactor that redirects its modification specificity from glutamate to serine^20,21^, and enables *trans*-PARylation of core histones, we examined how HPF1 affects PARP1-driven nucleosome disassembly. We adopted the previously reported substoichiometric concentration of HPF1 in our assay to reflect its physiological levels in cells^79^. In the presence of HPF1, we observed a decrease in the 10N30bio nucleosome band intensity concomitant with the formation of higher-order chromatin species, consistent with nucleosome *trans*-PARylation (Fig. 3A). These results demonstrated that a catalytic amount of HPF1 can efficiently promote PARP1-catalyzed *trans*-PARylation of chromatin. Notably, the hexasome species formed at early time points and disappeared at later time points, suggesting that these intermediates also underwent *trans*-PARylation (Fig. 3A). Importantly, we did not observe a substantial increase in free 10N30bio DNA even in the presence of excess quenching DNA, indicating that the majority of PARylated nucleosomes remained intact. Consistent with this observation, treatment of PARylated chromatins with poly(ADP-ribose) glycohydrolase (PARG) revealed distinct bands corresponding to mono-ADP ribosylated substrates (Fig. 3A).

**Figure 3.**
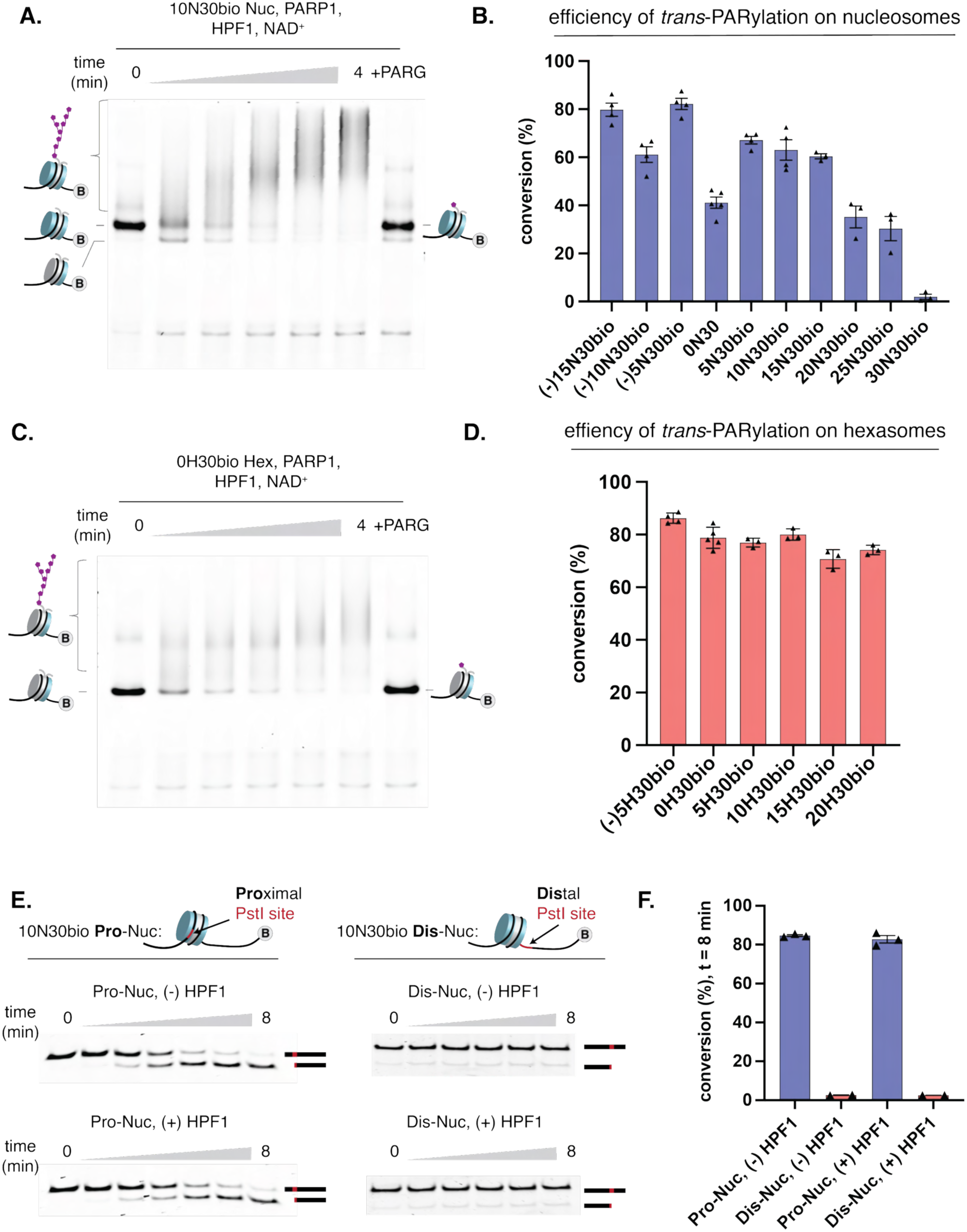
PARP1/HPF1 promotes nucleosome *trans*-PARylation while inducing nucleosome disassembly. (A) A substoichiometric amount of HPF1 facilitates *trans*-PARylation of the 10N30bio nucleosome to generate intact PARylated chromatin species that can be converted back to nucleosome structures upon treatment with PARG. Gel was stained with SYBR™ Gold to visualize total DNA. (B) Effect of linker length on nucleosome *trans*-PARylation. (C) A substoichiometric amount of HPF1 facilitates *trans*-PARylation of the 0H30 hexasome to generate intact PARylated chromatin species that can be converted back to hexasome structures by treatment with PARG. Gel was stained with SYBR™ Gold to visualize total DNA. (D) Effect of linker length on hexasome *trans*-PARylation. (E) Time-course analysis of chromatin accessibility during PARP1-mediated remodeling of 10N30bio Pro-Nuc and 10N30bio Dis-Nuc substrates in the presence and absence of HPF1, measured by a restriction enzyme accessibility assay (REAA). Reactions were initiated with PARP1, and aliquots were collected at the indicated time points and quenched with olaparib (12.5 μM). PstI was then added and incubated for 1 hour at 30 °C to digest accessible restriction sites. Following proteinase K treatment, the digested DNA was analyzed to assess restriction site accessibility. Gel was stained with SYBR™ Gold to visualize total DNA. (F) Quantification of disassembled nucleosomes from Fig. 3E at t = 8 minutes. HPF1, Histone PARylation factor 1; 0H30bio, a synthetic hexasome assembled on a 601-derived high-positioning sequence for hexasome (H) with 0 bp linker on one side and biotinylated 30 bp linker on the other side; 10N30bio Pro-Nuc: 10N30bio Nucleosome with a PstI site located proximal to the 10 bp linker; 10N30bio Dis-Nuc: 10N30bio Nucleosome with a PstI site located distal to the 10 bp linker. Chromatin remodeling reaction conditions: nucleosome (0.1 μM), PARP1 (0.2 μM), HPF1 (0.01 μM), NAD+ (0.5 mM).

Having established conditions to track chromatin state during *trans*-PARylation, we sought to assess how linker length affects chromatin PARylation (Figs. 3B, S3A-F). Similar to PARP1-mediated chromatin remodeling (Fig. 1E), the efficiency of nucleosome *trans*-PARylation is highly dependent on the length of the overhang DNA, with optimal PARylation occurring when linker lengths range from 5 to 15 base pairs (Fig. 3B). As the linker length approaches 30 base pairs, *trans*-PARylation efficiency drops significantly, consistent with the model in which PARP1 engages the DNA ends to promote histone *trans*-PARylation (Fig. 3B). We also observed elevated *trans*-PARylation upon truncation of DNA into the nucleosome core, following the same trend of nucleosome disassembly (Figs. 1E *vs.* 3B). Like nucleosome *trans*-PARylation, a catalytic amount of HPF1 can efficiently catalyze the complete modification of hexasomes (Figs. 3C, D, S3G-I). Overall, hexasomes exhibit higher activity than the corresponding nucleosomes carrying a similar linker length (Figs. 3B vs. 3D). Notably, the dependence of hexasome activity on linker length was less pronounced than in the nucleosome, with progressively diminishing activity as the linker DNA length increased (Fig. 3D). Moreover, hexasome PARylation resulted in a negligible increase in free DNA, and treatment of the PARylated hexasomes with PARG regenerated hexasomal species, suggesting that PARylated hexasomes remained intact (Figs. 3C, S3G-I).

We next evaluated how *trans*-PARylation of histones affects nucleosome disassembly by monitoring nucleosomal DNA accessibility using the restriction enzyme accessibility assay^59,80^. This assay was performed on 10N30 Pro-Nuc and Dis-Nuc, which carry a PstI site positioned proximal or distal to the 10-bp linker side (Fig. 3E). In the absence of HPF1, both Pro-Nuc and Dis-Nuc underwent nucleosome disassembly; however, only DNA from Pro-Nuc was efficiently digested by the enzyme, consistent with a mechanism in which PARP1 asymmetrically evicts the proximal histone dimer (Figs. S3J, 3E, and 3F). Similarly, although both Pro-Nuc and Dis-Nuc underwent HPF1/PARP1-mediated *trans*-PARylation, only the PstI site in Pro-Nuc was accessible, as indicated by efficient digestion (Figs. S3J, 3E, and 3F). Consistent with this model, enhanced accessibility of PstI sites proximal to the 10-bp linker was also observed in other nucleosomal and hexasomal substrates in the presence or absence of HPF1, suggesting that this mode of chromatin disassembly is a general feature under these conditions. (Figs. S3K and L).

It was previously reported that the *trans*-PARylation of histones primarily occurs at serine residues at the “KS” motifs, including H3S10, H3S28, and H2BS6^81,82^. As expected, alanine substitutions in the H3 tails (H3S10A/S28A) reduced nucleosome *trans*-PARylation, as reflected by the diminished level of slow mobility shift chromatin species (Fig. S3M). Notably, these mutations did not affect nucleosome disassembly (Fig. S3M). A similar trend was observed with transcriptionally active H3K9ac mark, which is known to block serine transPARylation (Fig. S3N). These data indicate that PARP1-driven nucleosome disassembly could be the dominant mechanism driving chromatin remodeling at genomic loci where the histone landscape suppresses serine PARylation.

Our systematic study demonstrates that sub-stoichiometric concentrations of HPF1 are sufficient to promote efficient *trans*-PARylation of chromatin. Furthermore, PARP1/HPF1-mediated nucleosome modification and disassembly can co-occur to generate stable PARylated hexasomes. These unique open chromatin forms may serve as bifunctional hubs for recruiting both DNA- and PAR-dependent factors.

### PARP activity drives chromatin accessibility with the direct involvement of subnucleosomes in the cellular context

Extensive studies using microirradiation-based assays support a model in which PARP activity-dependent chromatin remodeling promotes chromatin decompaction, facilitating the recruitment of DNA-binding proteins to damaged chromatin^12,27,29^; however, direct assays to measure chromatin accessibility remain lacking. Here, we developed protocols to dissect the contribution of PARP1 activity to chromatin accessibility in cells. We used well-established U2OS cells expressing PARP1-EGFP in a laser microirradiation assay, which revealed rapid recruitment of PARP1-EGFP to damage sites within seconds of irradiation (Fig. S4A)^11,12,83^. To directly assess chromatin accessibility, we repurposed DNA-footprinting-based sequencing technologies (fiber-seq and SAMOSA-seq) that use DNA methyltransferases to deposit N6-methyladenosine (m⁶A) marks on open chromatin, enabling direct visualization of accessible chromatin via antibody-based readout (Fig. 4A) ^84,85^. We refer to this new assay for open chromatin labeling and readout as SAMO-See. To capture chromatin accessibility at the early phase of the event, cells were fixed with formaldehyde within 1 minute of DNA damage induction (Fig. 4A). Notably, we observed the labeling signal along the laser path, suggesting enhanced chromatin accessibility at the damaged site (Figs. 4B,C). In the presence of Talazoparib, a potent PARP inhibitor, while PARP1 is still being recruited to the site, the m^6^A labeling signal significantly diminished concomitant with lower poly(ADP-ribose) (PAR) signal, consistent with the model that early chromatin accessibility is a PARP activity-dependent event (Figs. 4B,C). To exclude the possibility that adenine residues within PAR chains undergo m⁶A modification and thereby contribute to the observed m⁶A signal at damage sites, we treated fixed cells with PARG following DNA damage induction. The treatment abolished the PAR signal but had no effect on m⁶A labeling, indicating that m⁶A modification does not occur on the PAR chains (Figs. S4B,C). As expected, knockout of PARP1 abolished the SAMO-See signal and reduced PAR signal (Figs. 4D,E). Furthermore, PARP1-induced chromatin accessibility was also observed in HPF1 knockout U2OS cells, suggesting that chromatin accessibility at the early stage is at least in part independent of HPF1-mediated core histone *trans*-PARylation (Fig. S4D). A similar trend was observed with the neuroblastoma CHP212 cell line, suggesting the generality of the phenomenon (Fig. S4E). Consistent with enrichment observed in the in vitro pulldown assays (Figs. 2E, F) and previous studies^27,86^, we observed PARP-activity dependent recruitment of DNA factors, including Ku80, PRKDC, MRE11, and CHD4 (Figs. S4F-I).

**Figure 4.**
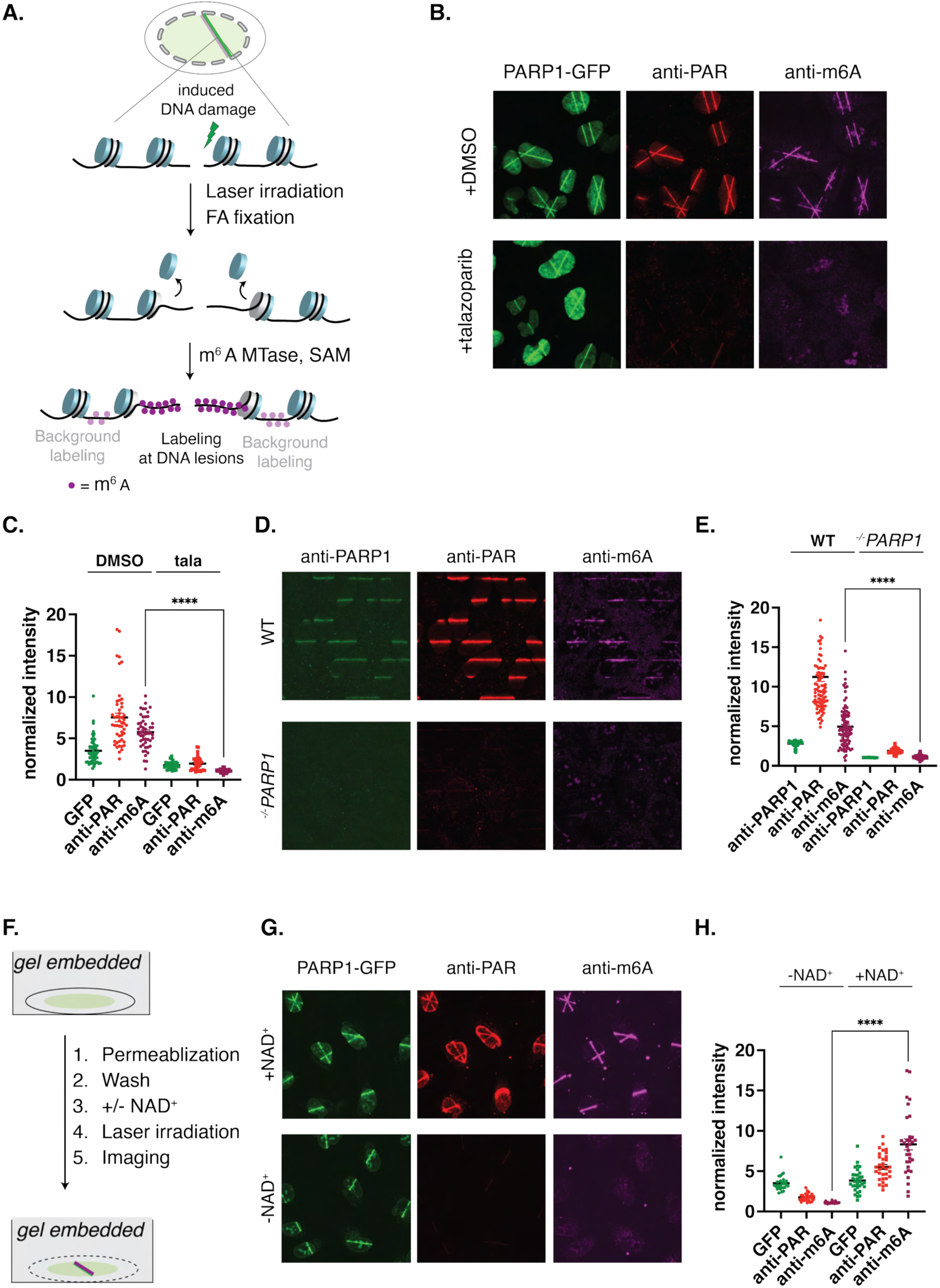
PARP activity directly promotes chromatin accessibility independent of ATP processes. (A) Schematic depicting a m^6^A-based DNA footprinting method for open chromatin labeling (SAMO-See) in a laser microirradiation assay. Following laser-induced DNA damage and fixation, permeabilized cells were treated with a non-specific m^6^A methyltransferase (MTase) to mark accessible chromatin with m^6^A, which was then detected using an antibody-based approach. (B) Chromatin accessibility at DNA lesions depends on PARP activity, as assessed by immunofluorescence for PAR and m^6^A. (C) Quantification of fluorescence intensity at sites of laser-induced DNA damage shown in Fig 4B. (D) Knockout of PARP1 reduces the PAR signal and abolishes the m^6^A signal. (E) Quantification of fluorescence intensity at sites of laser-induced DNA damage shown in Fig. 4D. (F) Schematic of nuclei embedding in gel, followed by metabolite removal and laser microirradiation. (G) NAD⁺-dependent PARP activity is required and sufficient to drive chromatin accessibility at DNA damage sites, as assessed by laser microirradiation of gel-embedded nuclei ± NAD⁺. (H) Quantification of fluorescence intensity at sites of laser-induced DNA damage shown in Fig 4G. Anti-PAR: antibody against Poly(ADP ribose).

To rule out the contribution of ATP-dependent processes, we developed an *ex vivo* nuclei imaging assay in which cells are permeabilized while embedded in the agarose gel and directly used for laser irradiation assay (Fig. 4F). This protocol enables the effective depletion of metabolites, including ATP, as measured by ATP-binding dyes, while maintaining PARP1-GFP recruitment activity upon activation of the laser light (Figs. S4J, K). Furthermore, PARP-mediated PARylation occurred upon the supplementation of NAD^+^ as measured by the appearance of the PAR signal (Fig. S4K). Importantly, we observed the SAMO-See signal along the laser path only upon addition of NAD^+^, suggesting that PARP activity directly induces chromatin accessibility (Figs. 4G,H). Furthermore, NAD^+^-dependent recruitment of Ku80 and PRKDC was observed in the *ex vivo* imaging assay (Figs. S4L-O). Interestingly, we did not observe CHD4 recruitment under similar conditions, suggesting that ATP-dependent processes may contribute to its recruitment to sites of damage (Fig. S4P). Together, these data suggest a direct effect of PARP1 activity on chromatin accessibility and the recruitment of several key DNA repair factors.

To investigate the contribution of subnucleosomal species such as hexasomes and tetrasomes, we adapted the in vitro nucleosome reassembly assay (Fig. 1C) for subnucleosome labeling of fixed nuclei (Fig. 5)^58,59^. To prevent histone dimer aggregation under our assay conditions, we established conditions for histone dimer transfer with a preformed histone dimer/DNA complex (Fig. 5A). Optimization for dimer complex formation was conducted on a short 33-bp dsDNA to reduce complex size and minimize multimeric oligomer formation (Fig. S5A). In vitro, this dimer complex efficiently transferred histone dimers to hexasomes, driving efficient nucleosome reassembly (Figs. S5B and 5B). In cellular assays, upon laser-induced DNA damage and mild fixation, chromatin reassembly was induced by treating permeabilized cells with HA-tagged dimer-DNA complexes (Fig. 5C). This strategy enables direct interrogation of subnucleosomal species within damaged chromatin by monitoring HA-tag incorporation. Importantly, we observed robust colocalization of the HA-tag signal with the laser-induced damage track (Figs. 5D,E), consistent with the local incorporation of histone dimers at sites of DNA damage. Notably, treatment with the PARP inhibitor Talazoparib markedly reduced the HA-tag signal, supporting a model in which the generation of subnucleosomal species is dependent on PARP activity (Figs. 5D,E). Importantly, treatment with PARG significantly reduces the PAR signal but does not affect dimer incorporation, suggesting that chromatin incorporation is at least in part independent of the PAR chain (Figs. S5C,D). Collectively, these data support a model in which PARP1 directly disassembles chromatin in response to DNA strand breaks, generating subnucleosomal species that promote chromatin accessibility and facilitate recruitment of repair factors.

**Figure 5.**
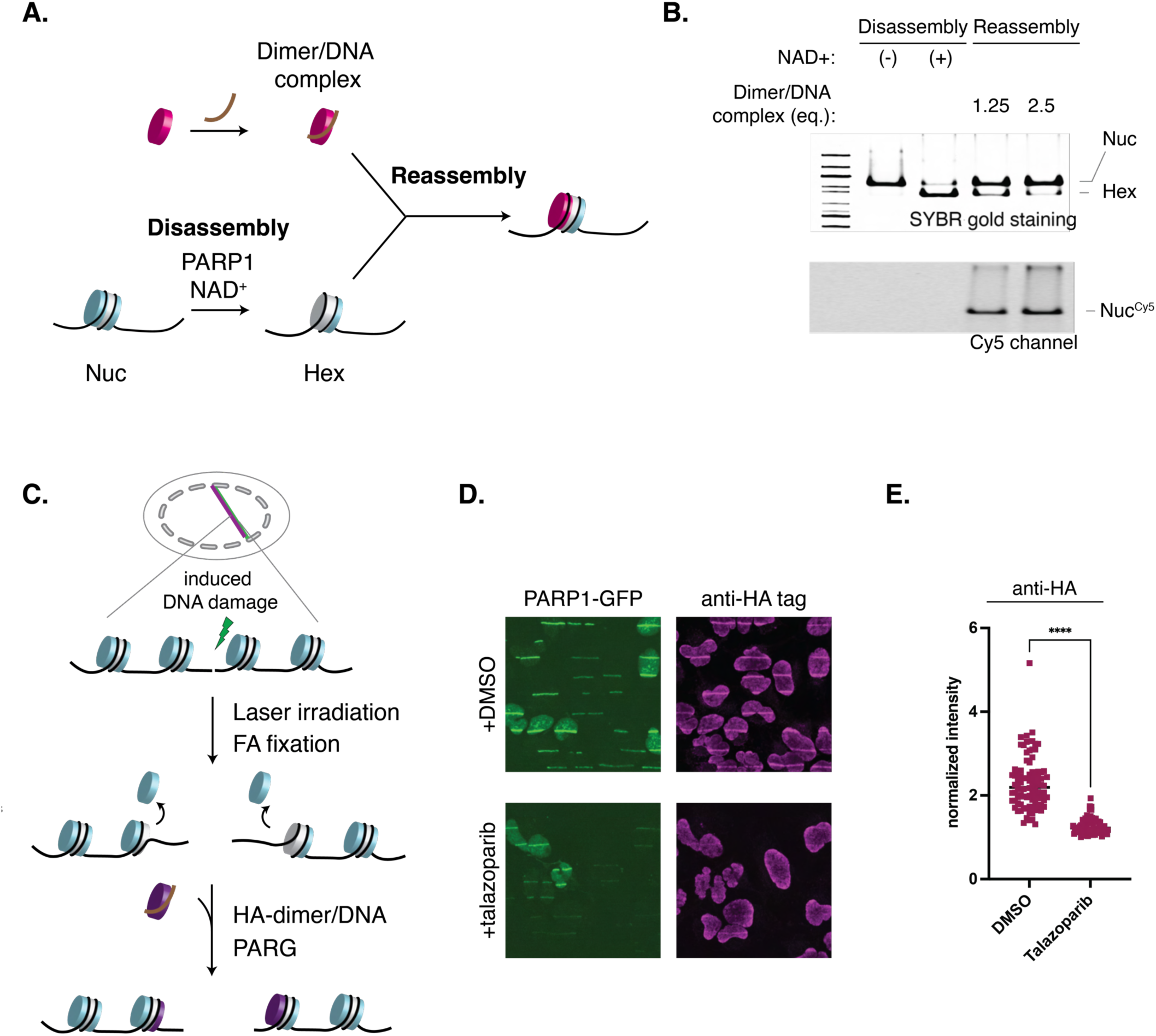
PARP-mediated nucleosome disassembly promotes the generation of subnucleosomes in a cellular context. (A) Strategy for nucleosome reassembly with a preformed dimer/DNA complex. (B) The preformed dimer/DNA complex efficiently transfers the histone dimer to the hexasome in a biochemical assay. (C) Schematic of in situ chromatin reassembly assay following laser microirradiation. Following laser-induced DNA damage and mild fixation, permeabilized cells were treated with the preformed HA-tagged dimer/DNA complex to allow nucleosome reassembly. Cells were then treated with PARG to remove the potential histone dimer/PAR complex. (D) Incorporation of HA-tagged dimer into DNA damage lesions is dependent on PARP activity. (E) Quantification of HA-tag fluorescence intensity at the laser-induced DNA damage sites shown in Fig 5D.

### C-terminal tail of H2A regulates PARP1-mediated nucleosome disassembly to promote DNA repair

To determine the histone domain that regulates PARP1-mediated nucleosome disassembly, we conducted a structure-activity relationship screening by performing the PARP1-induced nucleosome disassembly assay with a nucleosome library carrying mutations at various locations (A.V and H.T.D., unpublished data). Unexpectedly, we found that deletion of 10 amino acids at the C-terminal tail of H2A abolished the nucleosome disassembly (Fig. 6A) while not affecting the HPF1-dependent *trans*-PARylation (Fig. S6A). Previous works highlighted the role of the H2A C-terminus in nucleosome stability and regulation by the ISWI family of ATP-dependent chromatin remodelers^73,87,88^. While the original work by Längst and Schneider found the functional importance of the H2A C-terminus under stress conditions, the underlying molecular mechanisms remained unclear^88^. Our screening data suggest an unrecognized mechanism by which the protruding region of H2A facilitates PARP1-mediated chromatin disassembly at DNA lesions early in the repair process.

**Figure 6.**
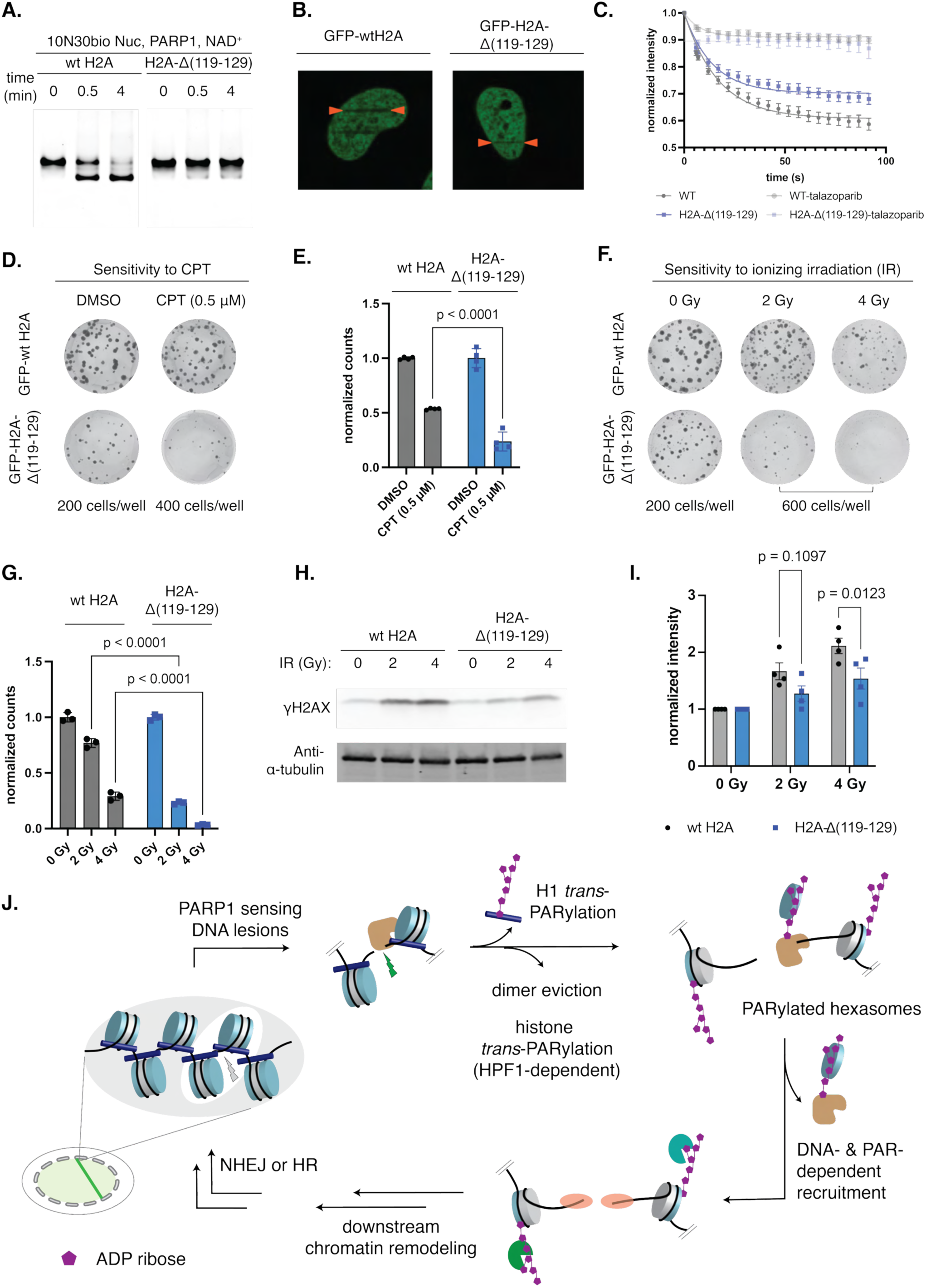
The H2A C-terminus regulates PARP1-mediated histone displacement to promote DNA repair. (A) Deletion of the C-terminal H2A tail abolished PAPR1-mediated nucleosome disassembly in a biochemical assay. Gel was stained with SYBR™ Gold to visualize total DNA. (B) Deletion of the H2A C-terminal tail reduces PARP1-mediated histone displacement in a two-photon microirradiation assay. Representative images at t = 60 s; ROIs indicated by lines. (C) Quantification of GFP signal at laser-induced DNA damage sites shows a greater reduction in U2OS cells expressing wild-type H2A compared with H2A-Δ(119–129). (D) Deletion of the H2A C-terminal tail sensitizes cells to the DNA-damaging agent camptothecin (CPT). (E) Colony numbers from Fig. 6D were quantified across four independent experiments. Statistical significance was determined using two-way ANOVA followed by Sidak’s multiple comparison test. (F) Deletion of the C-terminal H2A tail sensitizes cells to ionizing radiation. (G) Colony numbers from Fig. 6F were quantified across three independent experiments. Statistical significance was determined using two-way ANOVA followed by Sidak’s multiple comparison test. (H) Deletion of the H2A C-terminal tail potentially delays ATM-dependent DNA damage response signaling. Western blot analysis showed reduced levels of γH2AX in cells expressing truncated H2A compared with cells expressing wild-type H2A at 2 hours following ionizing irradiation. (I) Densitometric quantification was derived from four replicates across three independent experiments. Statistical significance was determined using two-way ANOVA followed by Sidak’s multiple-comparisons test. (J) Working model for NAD^+^ dependent chromatin remodeling activity of PARP enzymes to generate PARylated hexasome intermediates that can facilitate both DNA- and PAR-dependent recruitment to promote DNA repair.

To study the effect of the histone H2A tail on PARP1 activity in the context of DNA repair, we generated stable U2OS cell lines expressing GFP-wtH2A and GFP-H2A-Δ(119-129), respectively. To evaluate histone eviction, we adopted a previously established assay to measure histone displacement by measuring the loss of GFP signal after DNA damage induced by two-photon laser irradiation (Fig. 6B). We optimized the laser settings to minimize direct GFP photobleaching and confirmed that GFP signal loss is PARP activity-dependent, as no signal loss was observed upon treatment with a PARP inhibitor (Fig. S6B). Consistent with the trend observed in our biochemical assays (Fig. 6A), U2OS cells expressing truncated GFP-H2A exhibited reduced loss of GFP signal relative to wild-type H2A (Figs. 6B,C, S6C). A similar trend was observed in *^−/-^HPF1* U2OS cell lines, suggesting that the decrease in histone displacement observed in truncated H2A expressing cells is, at least in part, independent of HPF1 (Fig. S6D). To assess the functional implications of the histone H2A tail in DNA repair, we evaluated cells’ growth under different conditions using colony-forming assays. Upon exposure to DNA strand break-inducing treatments, including camptothecin (CPT) and ionizing irradiation, U2OS cells expressing truncated H2A exhibited increased sensitivity to DNA damage relative to cells expressing wild-type H2A (Figs. 6D-G). The reduced γH2AX signal, a key DNA damage marker, in truncated H2A-expressing cells suggests a delay in the downstream ATM-dependent DNA repair signaling (Figs. 6H, I). Consistent with a model in which the H2A C-terminal tail modulates PARP1 activity, truncation of H2A sensitized cells to PARP inhibitor treatment both in the presence and absence of CPT (Figs. S6E-H). Together, these data suggest a critical role for the H2A C-terminus in regulating PARP1-mediated nucleosome disassembly, which directly impacts DNA repair.

## Discussion

In this work, we discovered a third role of PARP1, in which PARP1 mediates the NAD+-dependent disassembly of nucleosomes upon activation by single-stranded and double-stranded breaks. In the absence of HPF1, we observed the automodification of PARP1, while no obvious modification occurred to the core histones, suggesting that PARylated PARP1, but not the core histones, plays a major role in nucleosome disassembly. Furthermore, PARG-catalyzed degradation of PAR chains on PARP1 restores the nucleosome structure, suggesting the association of PAR chains with the histone dimer. Based on this evidence, we propose that autoPARylation of PARP1 facilitates the removal of the histone dimer by promoting the interaction between negatively charged PAR chains and basic histone proteins (Fig. 6J). This unexpected nucleosome disassembly activity of PARP1 extends the previous models developed by Althaus and Luger, which posit that automodification of PARP1 can induce “histone shuttling” and serve as a histone chaperone^74,89^. Although the possibility of PARP1-induced disassembly of nucleosomes had been investigated previously, it may have been overlooked due to substrate designs that potentially interfered with this specific PARP1 activity^89^. Interestingly, under the same conditions, we do not observe nucleosome disassembly with PARP2, a closely related PARP member that shares many similarities in activity. This unique function of PARP1 suggests the existence of an unknown PARP1-specific domain that is required for inducing nucleosome disassembly.

Building on Pascal’s model of the PARP1-dsDNA complex, we designed our synthetic chromatin substrates to explore the effect of DNA lengths on nucleosome remodeling and modification^17^. We observed maximal activity with nucleosome substrates carrying 10-15 bp overhangs, consistent with the estimated 45 Å distance from the ART domain to the DNA end predicted by their model. Interestingly, we observed increases in both nucleosome remodeling activity and histone *trans*-modification upon truncation of the nucleosomal core. Truncation of the nucleosomal DNA on one side will reduce interaction between the histone dimer and nucleosomal DNA on that side, lowering the barrier to dimer eviction from the nucleosome. Consistently, we also observed higher nucleosome disassembly efficiency with native DNA sequences. Furthermore, the H2B and H3 tails in truncated nucleosomes become more conformationally accessible than those in canonical nucleosomes, providing a structural basis for increased HPF1-dependent *trans*-PARylation. In the same line, the removal of one histone dimer from the nucleosome to form the corresponding hexasomes also renders the histone H2B and H3 tails of hexasomes more accessible, consistent with our observation that hexasomes exhibit a higher *trans*-PARylation activity relative to intact nucleosomes. Previous work reported that active promoter H3K9ac and enhancer H3K27ac marks inhibit *trans*-PARylation^81,82^. Our in vitro data showed that mutation or modification at these sites did not block histone eviction, suggesting that PARP1-mediated nucleosome disassembly might play a primary role in facilitating DNA repair at these DNA-damage-susceptible sites. We further found that chromatosomes undergo disassembly, suggesting that PARP1-mediated nucleosome disassembly can disrupt higher-order chromatin architecture and thereby contributes to overall chromatin relaxation. Consistent with previous studies, H1 was robustly PARylated by PARP1, while core histones remained largely unmodified in the absence of HPF1^24^. These observations suggest that H1 dissociation may be driven by charge-based repulsion between its PAR chains and DNA. In contrast, the electrostatic interaction between the proximal dimer and automodified PARP1 might contribute to the eviction of core histone dimers ^74,89^.

One key feature of PARP1-mediated chromatin remodeling activity is its site-specificity, as it asymmetrically removes the histone dimer proximal to DNA lesions, thereby producing the corresponding oriented hexasomes. Eviction of the histone dimer would expose 35–50 base pairs of DNA at the damage sites, rendering the broken chromatin more accessible to DNA-binding factors. This nucleosome disassembly mechanism supports the DNA-dependent recruitment of chromatin effector proteins, including the NURD complex ATPase subunits CHD4, which were observed at damaged lesions, independent of PAR, by Timinszky and Huet^27^. In addition to CHD4, we observed that key DNA repair factors with DNA-binding activity upstream of both NHEJ and HR pathways, including Ku80/Ku70^90^ and MRE1^91^, preferentially bind disassembled chromatin rather than unmodified substrates in chromatin pulldown assays with nuclear extracts. Consistent with our biochemical assays, our DNA footprinting–based SAMO-See assay, together with ex vivo nuclei imaging, provides evidence that PARP1 directly drives chromatin accessibility independent of ATP-dependent remodeling. Beyond histone *trans*-PARylation, which promotes chromatin relaxation and accessibility^18,23–26^, our in situ chromatin reassembly assays demonstrate that PARP1-dependent nucleosome disassembly generates subnucleosomal intermediates that further enhance chromatin accessibility. Importantly, in our biochemical assays, *trans*-PARylated hexasomes remain intact; thus, they can also serve as binding hubs that recruit downstream PAR-dependent effector proteins. The hexasome intermediate mechanism can explain the enrichment of both PAR-dependent and DNA-dependent recruitment of repair factors at DNA lesions (Fig. 6J).

Although our model suggests direct binding of the growing PAR chain to the histone dimer, our data also indicate that the H2A C-terminus functions as a regulatory motif that governs this unique chromatin-remodeling activity, potentially controlling PARP1 orientation on the nucleosome. Deletion of this H2A domain abolished PARP1-mediated nucleosome disassembly in vitro and markedly reduced histone displacement from cellular chromatin. Notably, cells harboring the truncated H2A exhibited heightened sensitivity to both DNA strand break-inducing agents and PARP inhibitors, supporting the role of PARP1-mediated nucleosome disassembly in DNA damage response. A previous study by Längst and Schneider highlighted the unresolved mechanism of the specific effects of H2A truncation on the stress response^88^. Our work provides both in vitro and cellular evidence supporting direct involvement of the H2A C-terminus in chromatin regulation via the PARP1 axis, explaining its role in DNA repair.

In conclusion, we uncover a previously uncharacterized chromatin-remodeling activity of PARP1 that drives NAD^+^-dependent nucleosome disassembly upon activation by DNA strand breaks. These findings provide a mechanistic framework for earlier observations of PARP-mediated chromatin relaxation, histone displacement, and recruitment of repair factors to DNA damage sites through both DNA- and PAR-dependent mechanisms. Deletion of the H2A C-terminus markedly impairs PARP-dependent histone displacement in both in vitro and cellular assays and sensitizes cells to DNA damage, establishing PARP-dependent nucleosome disassembly as a functionally relevant mechanism in DNA repair. Notably, several oncohistone mutations have been identified in this H2A domain^73^, underscoring its potential relevance to disease-associated chromatin dysfunction. More broadly, this study provides the first direct evidence linking subnucleosomes to the DNA damage response, setting the stage for future investigations into their functional roles.

### Limitation of the study

Although our biochemical analyses demonstrate that PARP1 can promote histone dimer eviction to generate asymmetric hexasomes, the precise structural intermediates formed at DNA breaks in cells remain to be determined. Future studies using high-resolution chromatin mapping will help define the structure of these subnucleosomal species in vivo.

---

**Figure S1.**
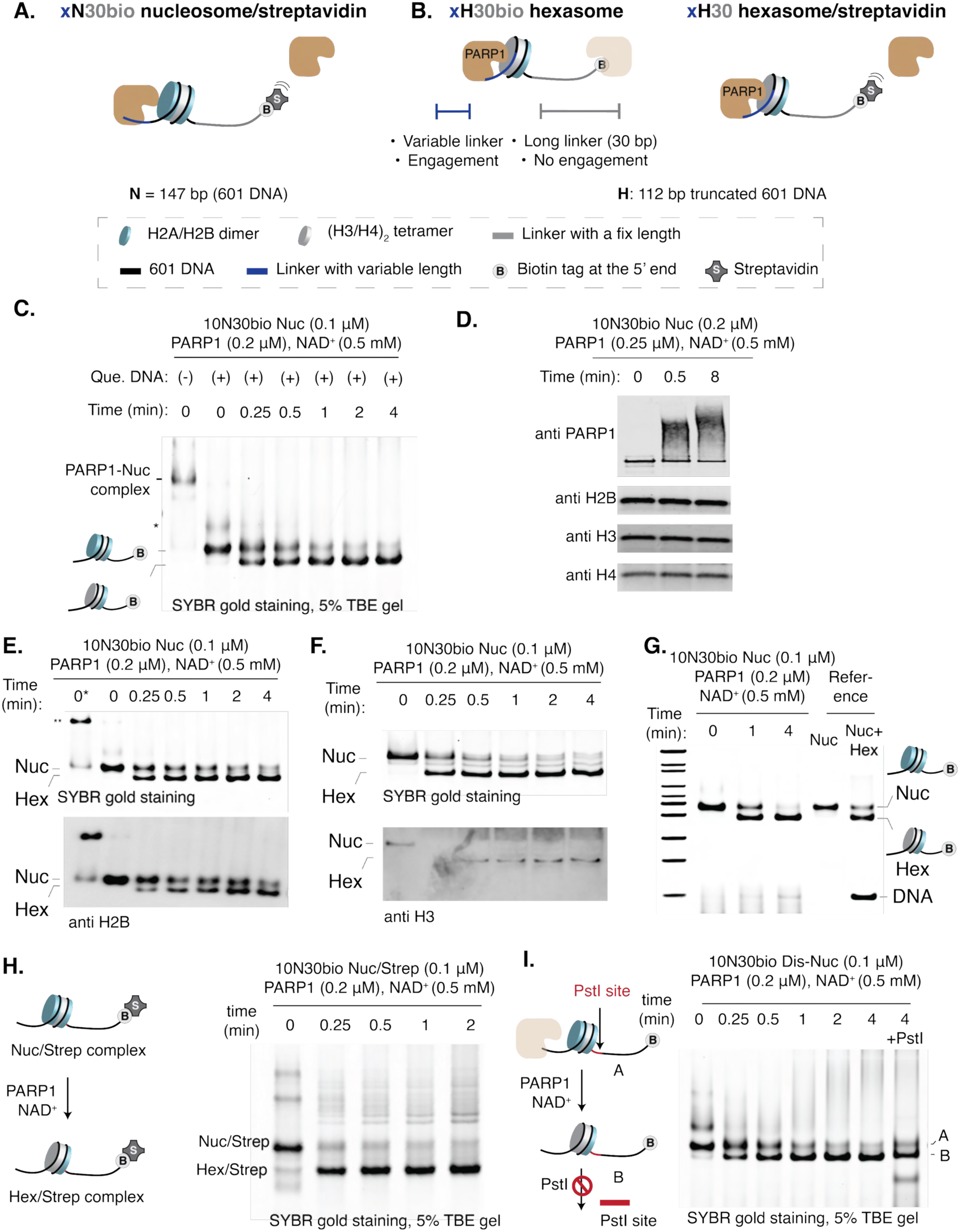
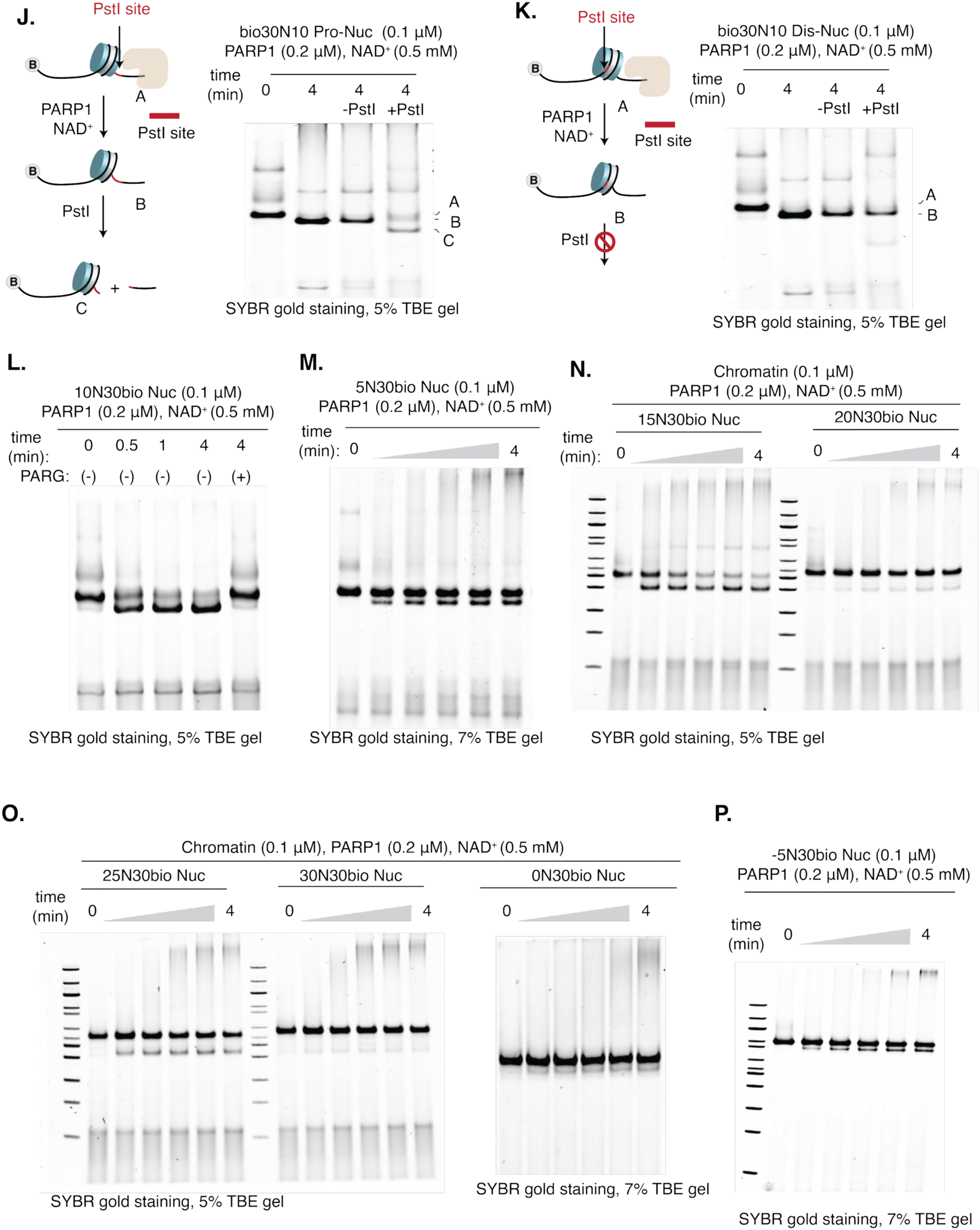
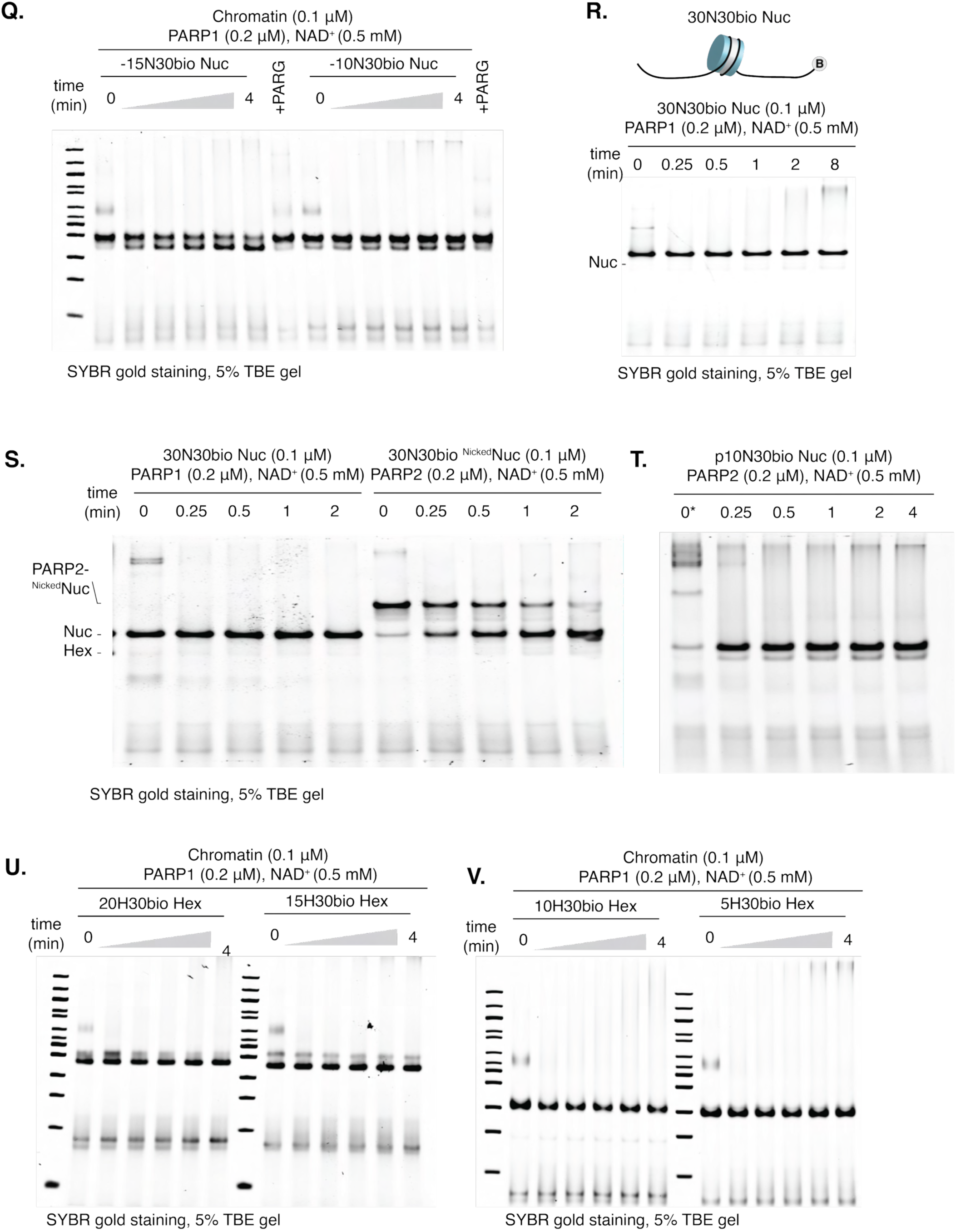
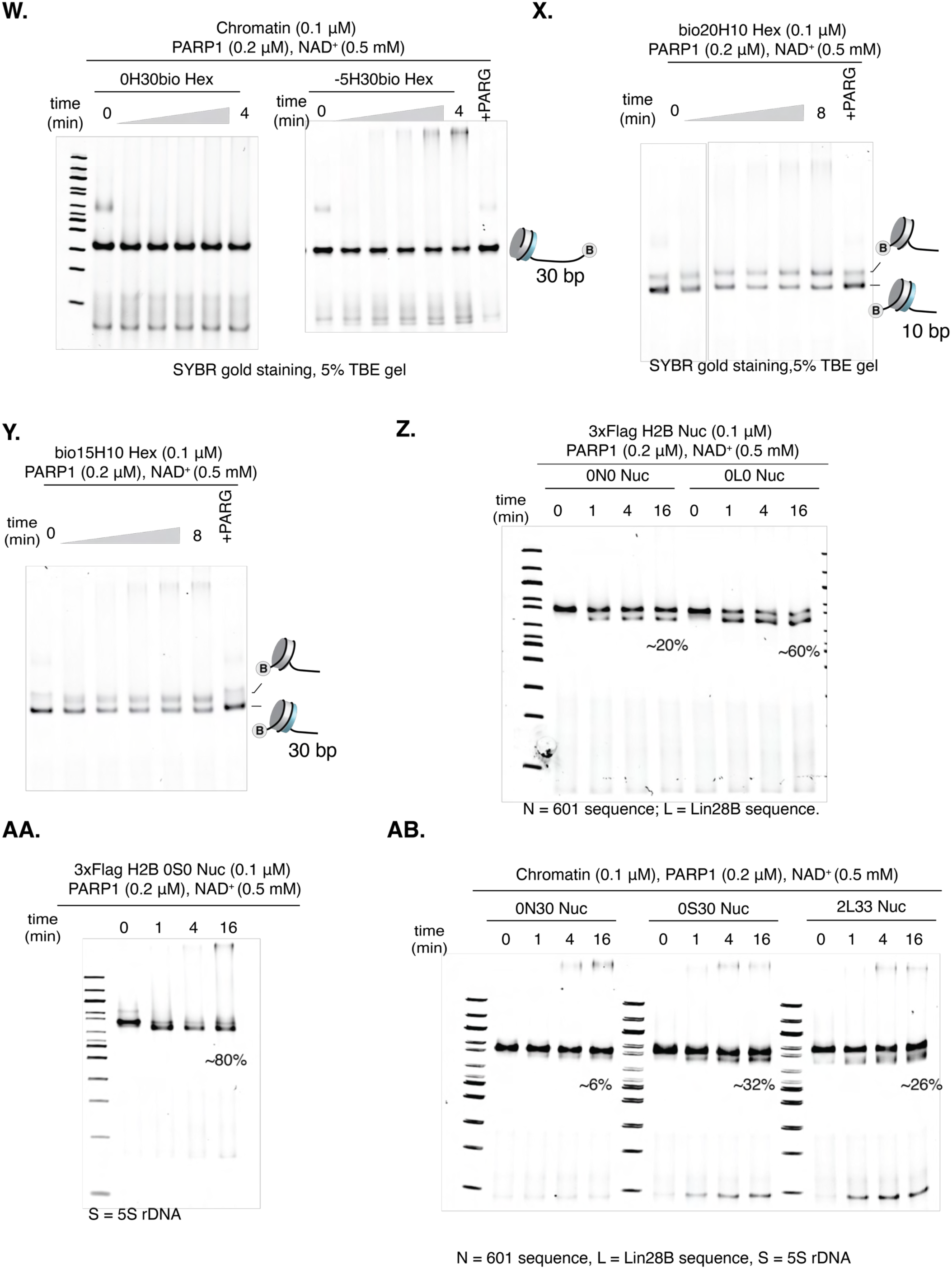
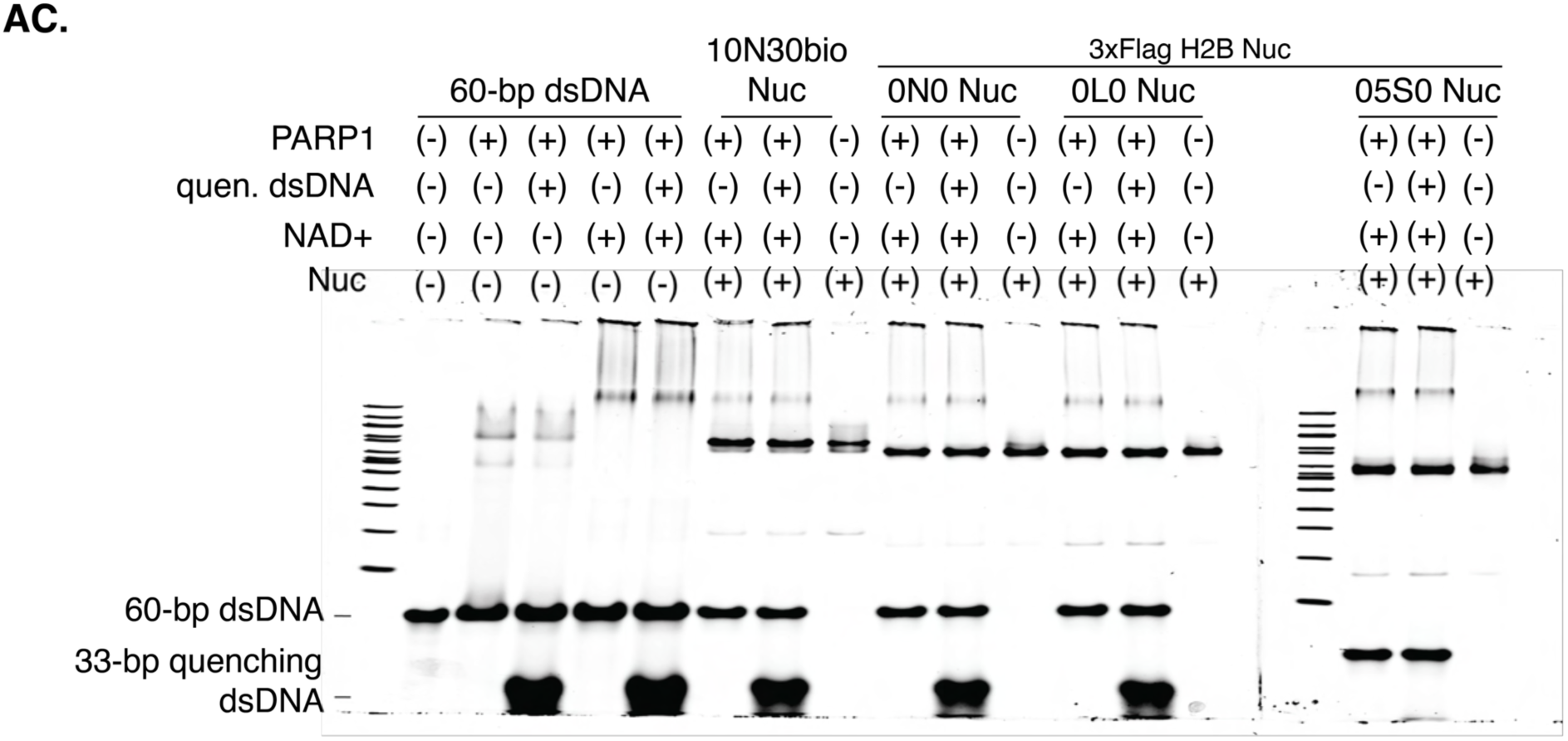
PARP1 asymmetrically disassembles chromatin. (A) Design of nucleosome substrates with a streptavidin blockage. (B) Design of hexasome substrates with and without a streptavidin blockage. (C) 60 bp dsDNA (quenching DNA) efficiently sequesters PARP1 from chromatin while preserving chromatin integrity, enabling tracking of chromatin structural changes during the reaction. (*) indicates a species tentatively assigned as a PARP1/60bp-dsDNA complex. (D) PARP1 undergoes *trans*-PARylation, whereas core histones remain unmodified, as determined by western blot analysis. (E,F) Fast-migrating chromatin species contain both H2B (E) and H3 (F) as determined by native gel electrophoresis followed by western blot analysis. In S1E: (*) indicates no quenching DNA; (**) indicates the putative PARP1-nucleosome complex. (G) Fast-migrating chromatin species exhibits mobility comparable to that of a reference hexasome. (H) Generation of fast-migrating chromatin species with chromatin containing streptavidin blockage. (I) The PstI site installed on the distal side to the 10 bp linker remained inaccessible. (J,K) PARP1-mediated nucleosome disassembly followed by restriction enzyme accessibility assay on bio30N10 substrates. (L) Disassembled nucleosome can be converted back to nucleosomes upon treatment with Poly (ADP-ribose) glycohydrolase (PARG). (M-Q) PARP1-mediated nucleosome disassembly as a function of the linker length. (R) The 30N30 nucleosome without a nicked site remains unchanged under similar conditions as in 1F. (S,T) PARP2 does not promote nucleosome disassembly under conditions comparable to those used for PARP1.(U-W) PARP1 does not disassemble hexasomes with the long 30bp linker positioned proximal to the dimer. (X,Y) PARP1 disassembles hexasomes carrying a 10 bp linker positioned proximal to the dimer. (Z,AA) Nucleosomes assembled on native DNA sequences (human LIN28B DNA or 5S rDNA) are more susceptible to disassembly than those assembled on 601 DNA. A 3×FLAG-tagged H2B was used to enhance the separation of zero-linker nucleosomes and the corresponding hexasomes on native gels. (AB) Nucleosomes assembled on native DNA sequences (human LIN28B DNA or 5S rDNA) are more susceptible to disassembly than those assembled on 601 DNA. The 0S30 nucleosome was tentatively assigned because it shows a mobility shift similar to that of the 0N30 nucleosome. 2L33 was assigned based on a Cryo-EM structure (PDB 8G8G, ref. 75). (AC) Pre-activated PARP1 does not disassemble nucleosomes. PARP1 was activated with dsDNA fragments and NAD+, followed by incubation with different nucleosomes in the presence and absence of quenching 33 bp-dsDNA.

**Figure S2.**
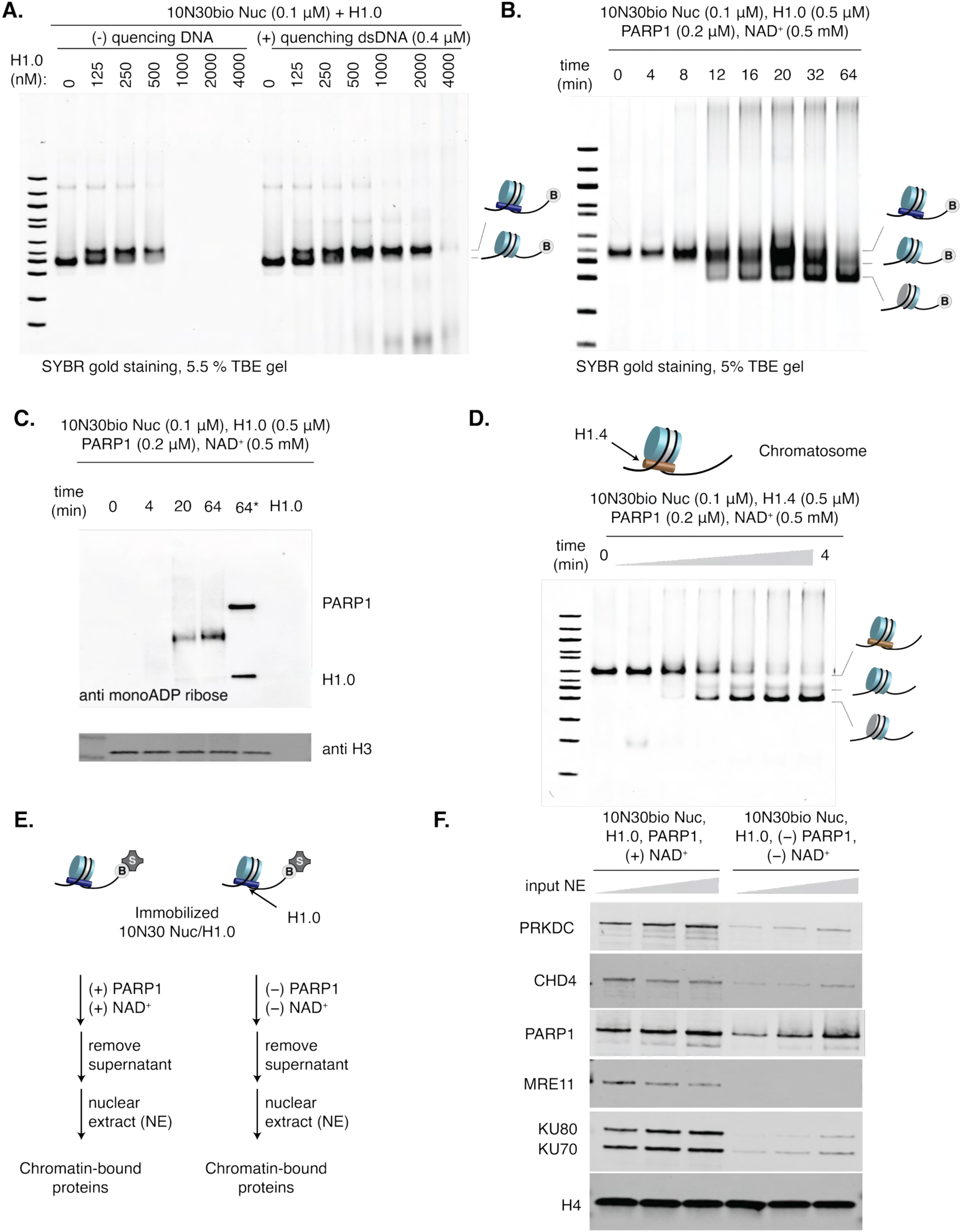
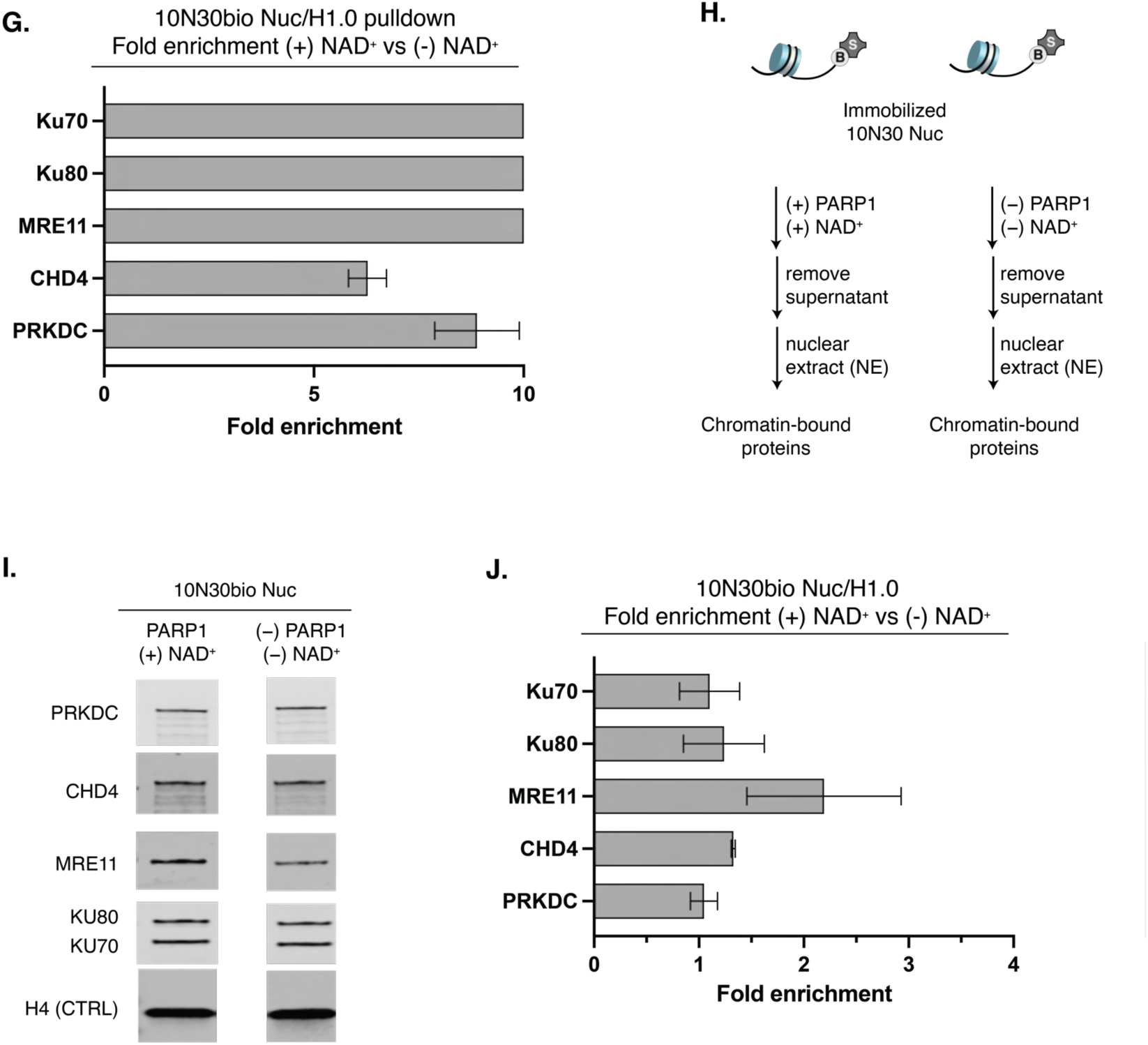
PARP1 disassembles chromatosomes to enhance chromatin accessibility to DNA repair factors. (A) Formation of H1.0/nucleosome complexes in the presence and absence of quenching 60 bp dsDNA. (B) Time course analysis of PARP1-mediated disassembly of nucleosome/H1.0 chromatosome complexes. (C) PARG treatment of remodeled chromatin (well 4) restored chromatosomes containing mono-ADP-ribosylated H1.0 (well 5), as assessed by Western blot. (D) PARP1 disassembles H1.4 chromatosome complexes. (E) Schematic of the chromatin pulldown assay with HeLa nuclear extract used to assess the effects of PARP1-driven chromatosome disassembly. In contrast to experiments in Fig 2D, PARP1 was excluded from the (-) NAD+ conditions. (F) PARP1-induced chromatosome remodeling promotes the recruitment of DNA repair factors. PARP1-induced chromatin remodeling enhances the binding of DNA repair factors, including Ku80, Ku70, PRKDC, MRE11, and CHD4. (G) Quantification of fold enrichment of repair factors in the presence of NAD⁺ relative to reactions lacking NAD⁺ in the experiment shown in Fig. S2F, under the lowest nuclear extract (NE) input conditions. Error bars represent SEM from two independent replicates. Fold enrichment values greater than 10 were capped at 10 due to the dynamic range limitations of Western blot detection. (H) Schematic of the chromatin pulldown assay with HeLa nuclear extract used to assess the effects of PARP1-driven nucleosome disassembly. In contrast to experiments in Fig 2D and Fig. S2F, H1.0 was excluded from the experiment. (I) The small enrichment of the remodeled nucleosome is observed for MRE11 and CHD4 but not for Ku80, Ku70, or PRKDC. (J) Quantification of fold enrichment of repair factors in the presence of NAD⁺ relative to reactions lacking NAD⁺ in the experiment shown in Fig. S2I. Error bars represent SEM from two independent replicates.

**Figure S3.**
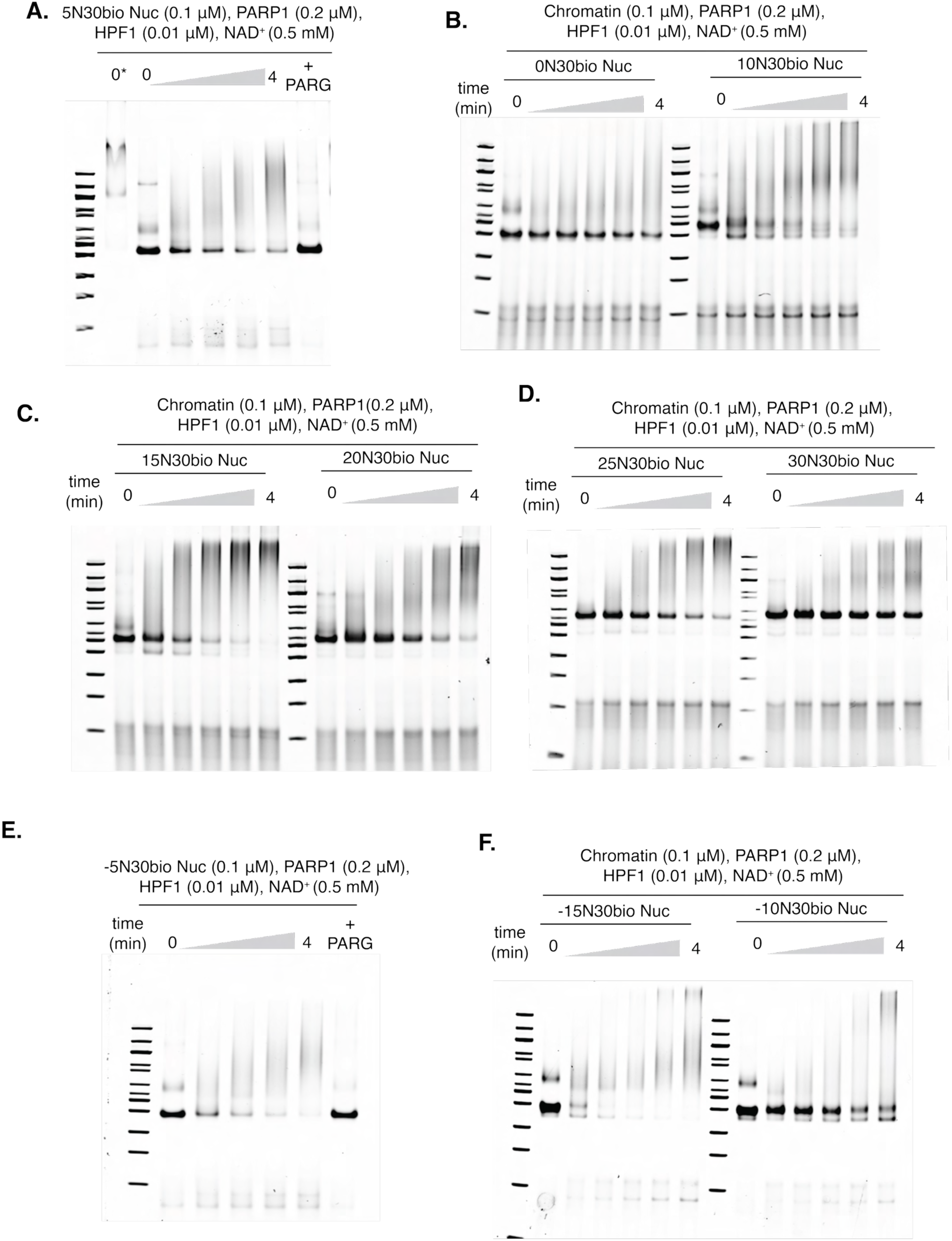
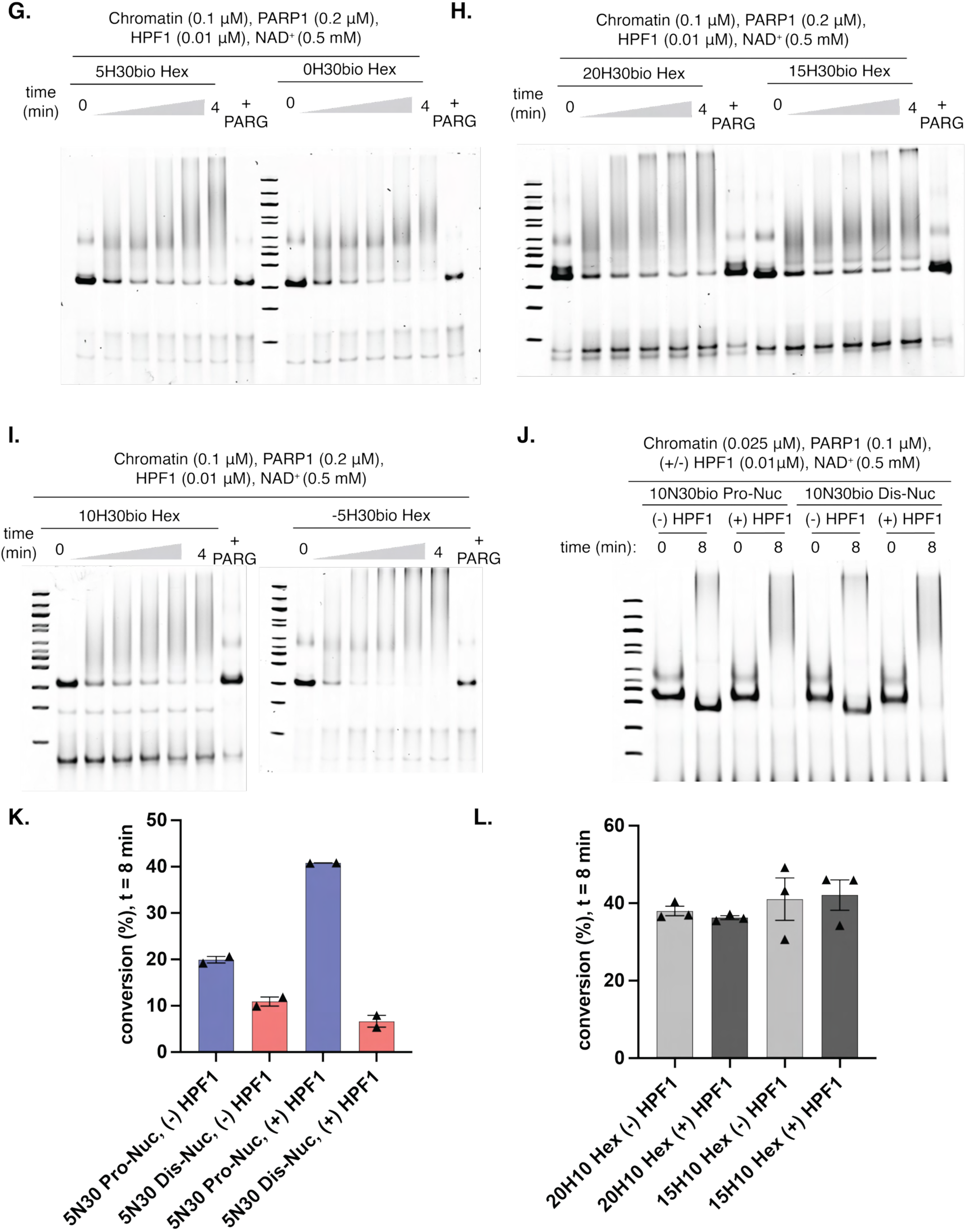
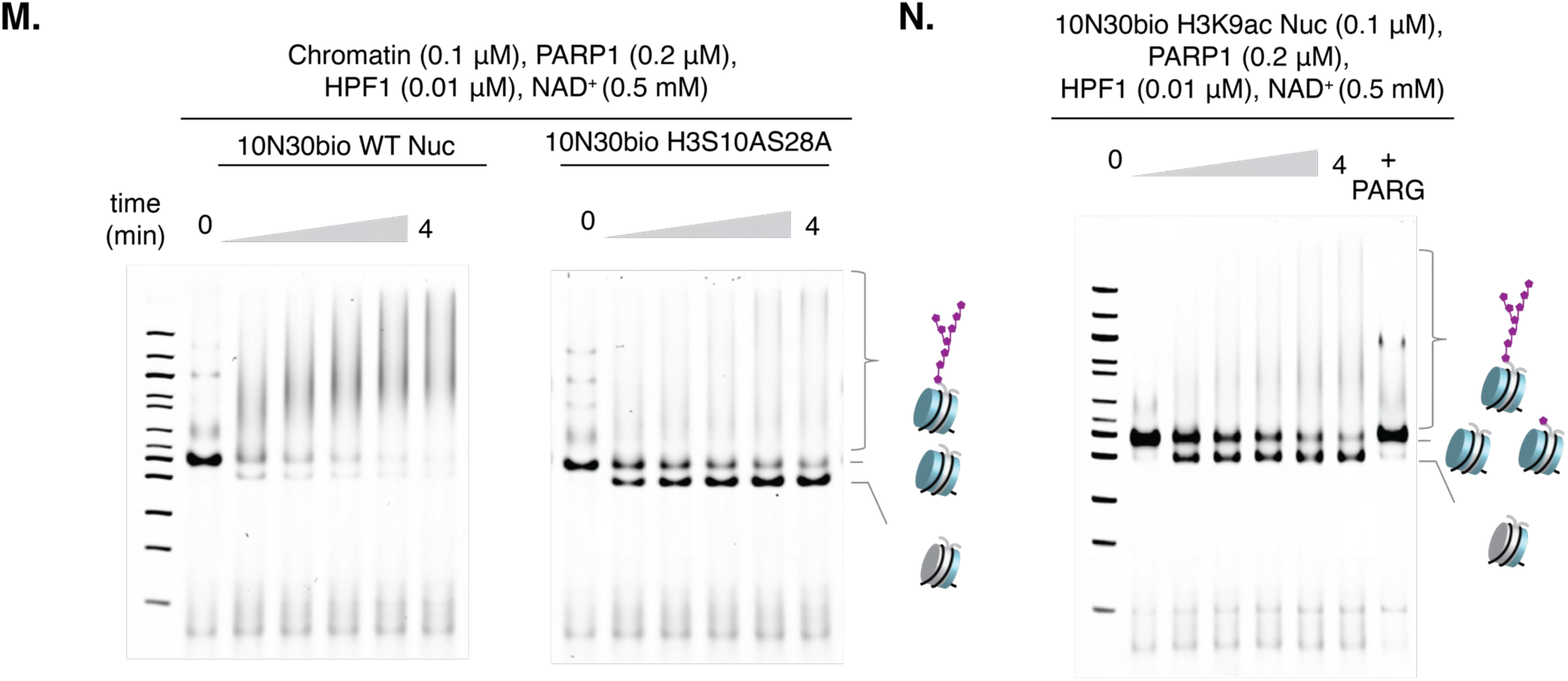
PARP1/HPF1 promotes both histone *trans*-PARylation and nucleosome disassembly. (A-F) Substoichiometric amount of HPF1 facilitates *trans*-PARylation of nucleosomes to generate PARylated chromatin species (G-I) Substoichiometric amount of HPF1 facilitates *trans*-PARylation of hexasomes to generate PARylated chromatin species that can be converted back to nucleosome structures upon treatment with Poly (ADP-ribose) glycohydrolase (PARG). (J) 10N30bio Pro- and 10N30bio Dis-nucleosomes underwent disassembly and modification in the absence and presence of HPF1. Aliquots were taken at t = 8 minutes. (K) Chromatin accessibility (t = 8 minutes) in PARP1-mediated chromatin remodeling of 5N30bio Pro-Nuc and 5N30 Dis-Nuc in the absence and presence of HPF1 as assessed by REAA. While distal sides remain inaccessible, the proximal sides become more accessible upon triggering PARP1 activity. A significant increase in accessibility to 5N30 Pro-Nuc was observed in the presence of HPF1. (L) Chromatin accessibility (t = 8 minutes) in PARP1-mediated chromatin remodeling of bio20H10 and bio15H10 hexasomes in the absence and presence of HPF1 as assessed by REAA. These hexasomes were shown to undergo disassembly in Figs. 1 X,Y. (M) Alanine mutation at acceptor sites (H3S10AS28A) abolishes *trans*-PARylation but not nucleosome disassembly as monitored in EMSA. (N) Acetylation of H3K9ac reduces *trans*-PARylation but did not affect nucleosome disassembly as monitored in EMSA.

**Figure S4.**
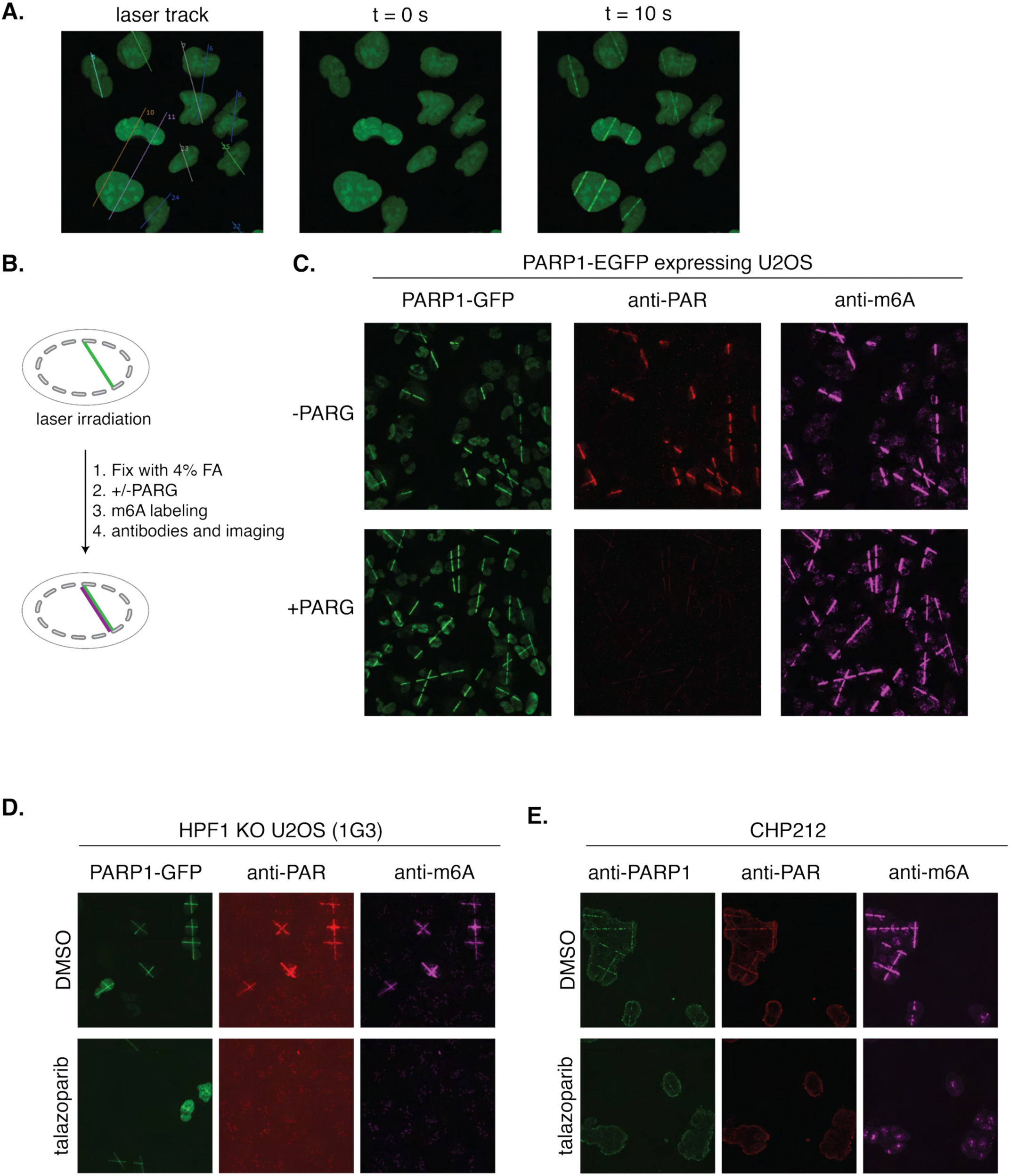
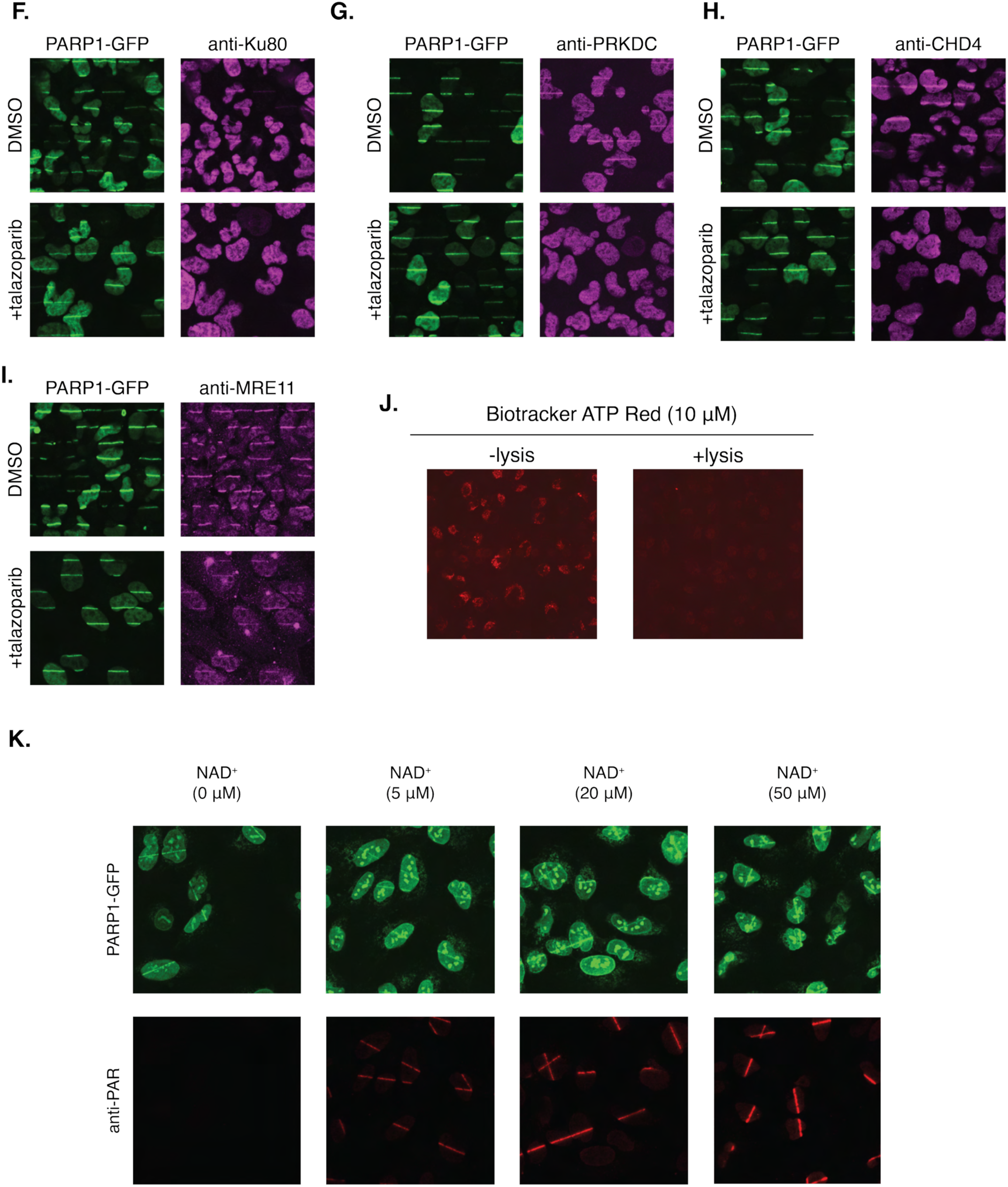
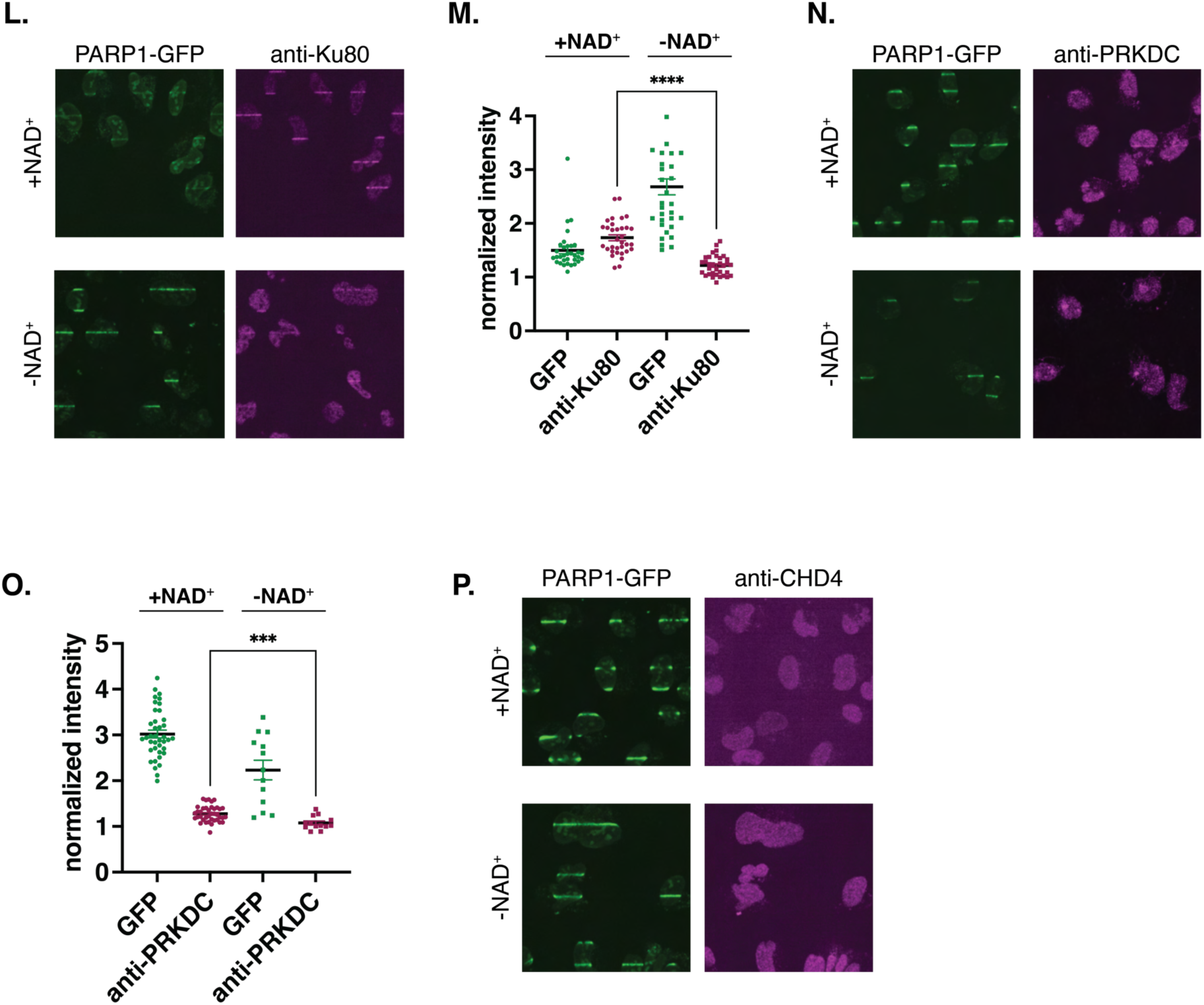
PARP activity is required and sufficient to induce chromatin accessibility and promote the recruitment of DNA repair factors to sites of damage independent of ATP processes. (A) Laser microirradiation reveals rapid recruitment of PARP1 to sites of laser-induced DNA damage. (B) Schematic of the experiment design used to exclude the potential contribution of direct m^6^A labeling on the PAR chains. PARG was added to degrade PAR prior to m6A labeling. (C) Chromatin accessibility at the laser track is independent of the PAR chain formation as the PARG treatment did not reduce m^6^A signal. (D) PARP activity-dependent chromatin accessibility was observed in HPF1 knockout U2OS cells. (E) PARP activity-dependent chromatin accessibility was observed in the neuroblastoma CHP212 cell lines. (F-I) PARP activity-dependent recruitment of DNA repair factors (Ku80, PRKDC, CHD4, and MRE11) to the laser track as assessed by immunofluorescence imaging. (J) ATP depletion was achieved following cell permeabilization and removal of metabolites from gel-embedded cells, as validated by ATP-binding dye imaging. (K) Under optimized permeabilizing and washing conditions, PARP1 remains competent for recruitment to the laser track and can be activated by the addition of NAD+. (L) NAD^+^ dependent chromatin remodeling activity of PARP enzymes facilitates the recruitment of Ku80 to damage lesions independent of ATP processes, as assessed by in-gel laser irradiation. (M) Quantification of fluorescence intensity at sites of laser-induced DNA damage shown in Fig. S4L. (N) NAD^+^ dependent chromatin remodeling activity of PARP enzymes facilitates the recruitment of PRKDC to damage lesions independent of ATP processes, as assessed by in-gel laser irradiation. (O) Quantification of fluorescence intensity at sites of laser-induced DNA damage shown in Fig. S4N. (P) CHD4 did not localize to laser-induced DNA damage tracks in gel-embedded nuclei microirradiation experiments.

**Figure S5.**
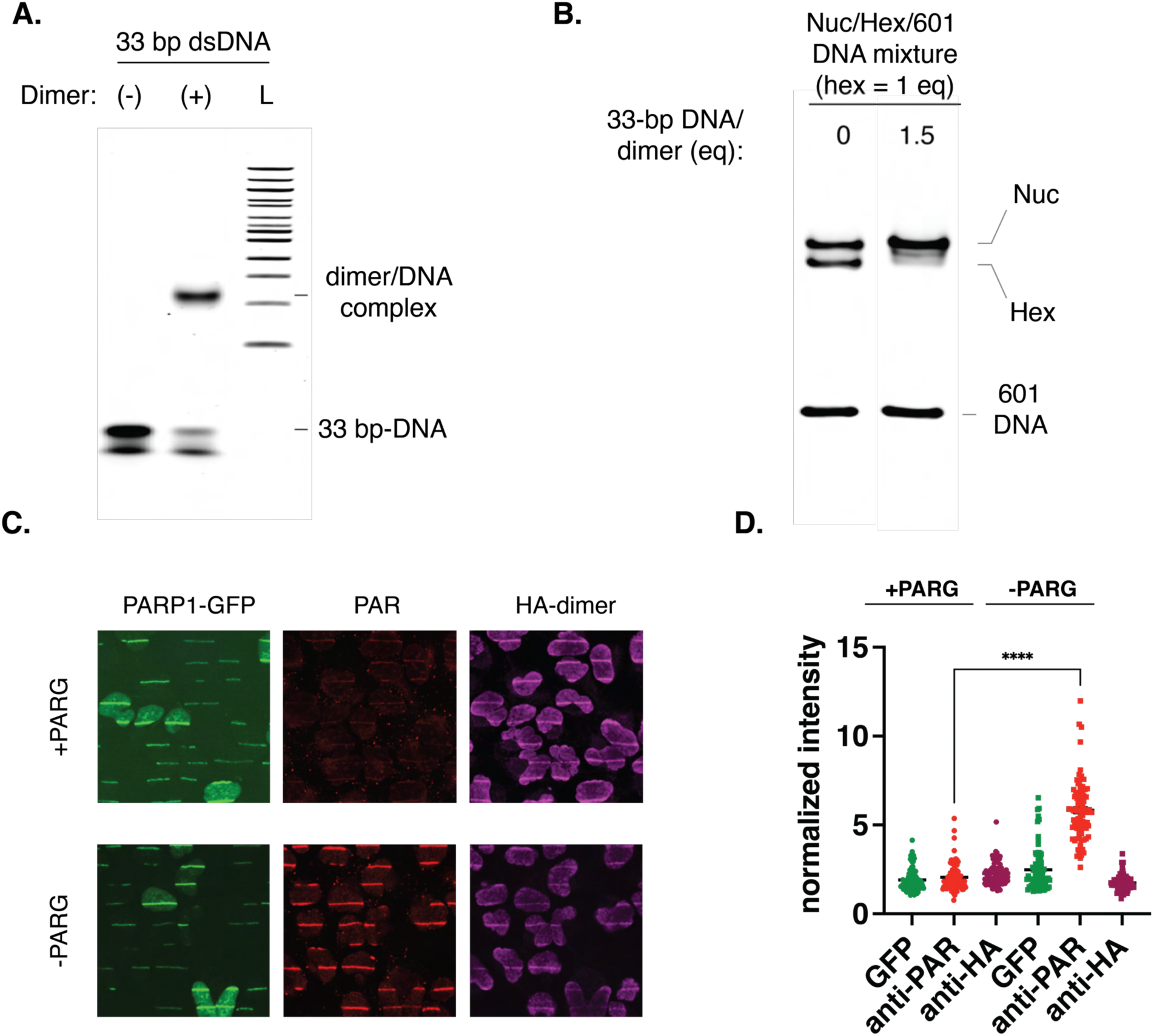
PARP-mediated histone displacement generates subnucleosomes in cellular assays. (A) Generation of a histone dimer/33bp-dsDNA complex as assessed by native gel electrophoresis. Gel was stained with SYBR™ Gold to visualize total DNA. (B) Dimer/DNA complex efficiently transfers the histone dimer to a hexasome in the presence of a nucleosome and the free 601 DNA in a biochemical assay. Gel was stained with SYBR™ Gold to visualize total DNA. (C) PARG treatment effectively abolished PAR signal but does not affect the HA signal. (D) Quantification of fluorescence intensity at sites of laser-induced DNA damage from experiments in Fig. S5C.

**Figure S6.**
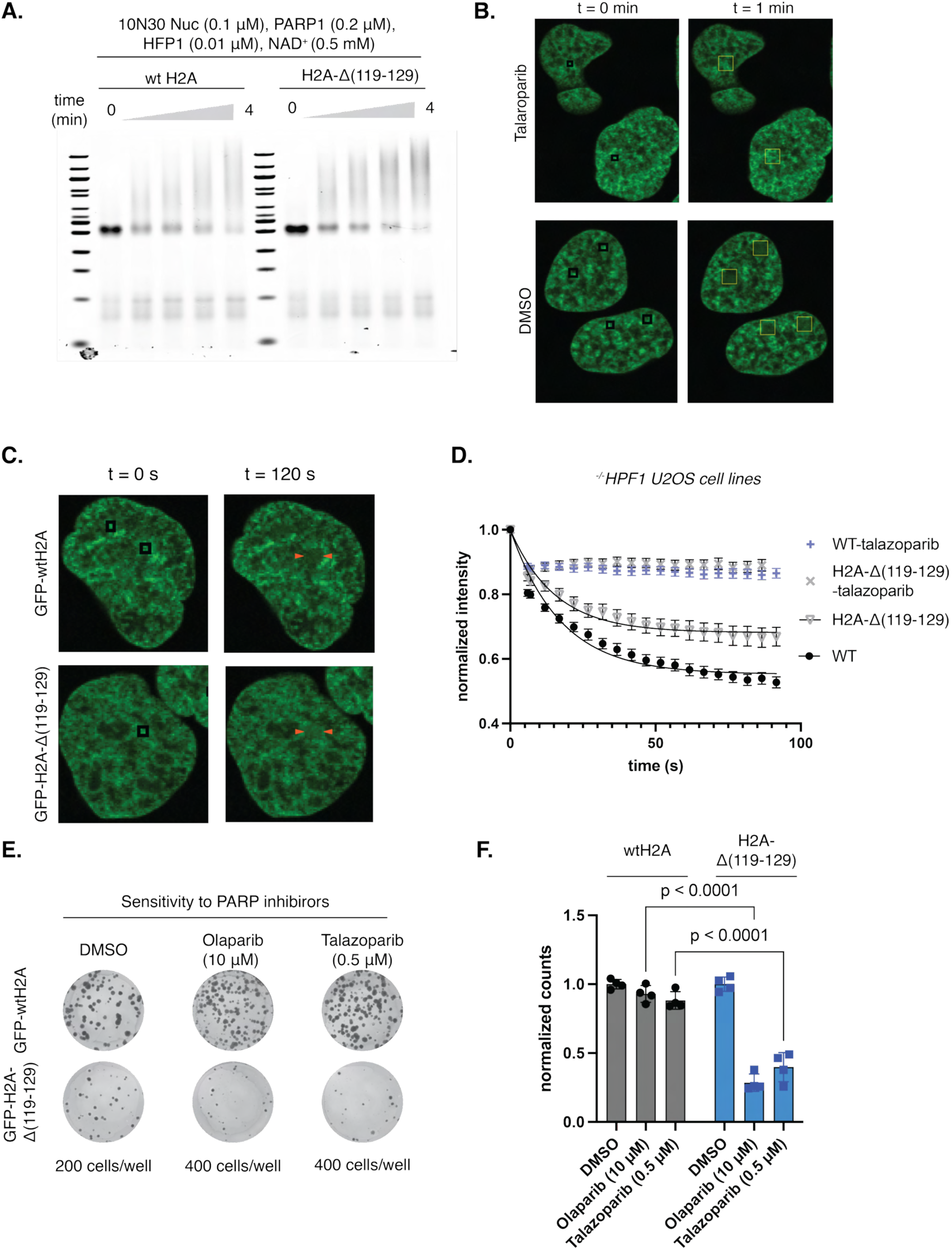
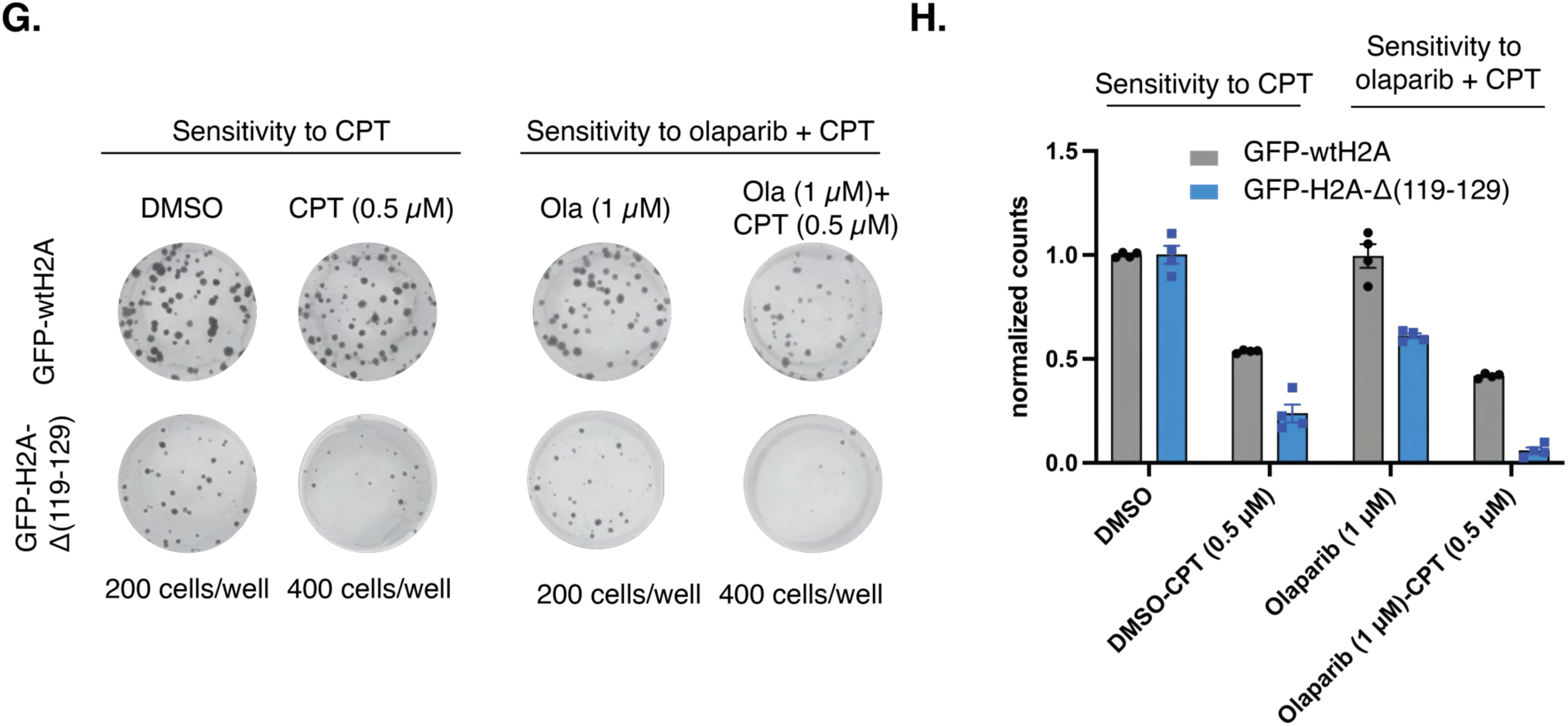
Deletion of the H2A C-terminus abolishes PARP1-mediated histone eviction in biochemical and cellular assays and impairs DNA repair. (A) Deletion of the C-terminal H2A tail abolishes PAPR1-mediated nucleosome disassembly but does not affect HPF1-mediated trans-PARylation. (B) Optimized two-photon laser irradiation settings that allow probing the PARP-dependent histone displacement. (C) Deletion of the C-terminal H2A tail reduces PAPR1-mediated histone displacement in a two-photon microirradiation assay. Representative images at t = 120 s; ROIs indicated by squares. (D) Quantification of GFP signal at laser-induced DNA damage sites shows a greater reduction in *^−/-^HPF1* U2OS cells expressing wild-type H2A compared with H2A-Δ(119–129). (E) Deletion of the H2A C-terminal tail sensitizes cells to the PARP1 inhibitors olaparib (10 µM) and talazoparib (0.5 µM). (F) Colony numbers from Fig. S6E were quantified from four independent experiments. Statistical significance was determined using two-way ANOVA followed by Sidak’s multiple comparison test. (G) Deletion of the H2A C-terminal tail sensitizes cells to the combination treatment of olaparib (1 µM) and CPT (0.5 µM). (H) Colony numbers from Fig. S6G were quantified across four independent experiments. Statistical significance was determined using two-way ANOVA followed by Sidak’s multiple-comparisons test. Quantified data from Fig. 6D are included for ease of comparison.

## Author Contributions

H.T.D. conceived the project and supervised the study. A.V. and H.T.D. performed biochemical experiments. A.P. and H.T.D. established the laser microirradiation assays. C.Z., B.T. and H.T.D. performed cellular assays. C.Z. and R.D. performed ionizing irradiation assays. S.B. and M.H. provided conceptual advice on biochemical experiments and contributed protein materials. A.S. provided conceptual advice on colony formation and ionizing irradiation assays. H.T.D. wrote the manuscript with input from all authors.

## Acknowledgments

We thank A. B. Taylor from the Cell and Tissue Imaging Center – Light Microscopy for advice on Fiji-based data processing, C. Deane for assistance with manuscript editing, and A. Z. Ansari for critical reading of the manuscript. We thank S. Miller and J. Quarterman from the Center for Advanced Genome Engineering (CAGE) for generating knockout cell lines, D. Vocelle from the Flow Cytometry and Cell Sorting Shared Resource for cell sorting, and P. Rodrigues from the Hartwell Center Sequencing Facility for synthesis of the hydrazide peptide. We thank R. Shrestha for assistance with chromatin pulldown experiments, O. Konar for assistance with biochemical assays of truncated H2A, and Y. Bhat for preparation of tagged histone dimers. We thank J. Teague (Halic laboratory) for providing protein materials. We thank A. Z. Ansari, R. Lee, T. Mittag, and S. Baker for strategic advice throughout the study. We also thank members of the SJ ChromaTeam (Ansari, Dao, Gibson, and Halic laboratories) for insightful discussions and the Chen, Ansari, and Lee laboratories for access to instrumentation.

## Supporting Information

### GENERAL METHODS

#### General information

Primers were purchased from Integrated DNA Technologies (Coralville, IA). DNA purification kits were purchased from QIAGEN (Valencia, CA) for DNA purification. Mutagenesis was conducted using the Inverse PCR method and Gibson assembly. DH5a and One Shot™ Stbl3™ Chemically Competent *E. coli* were generated in-house from cells purchased from NEB and Invitrogen. All plasmid sequences were verified by Sanger sequencing in-house or by Plasmid sequencing performed by Plasmidsaurus (San Francisco, CA). Size-exclusion chromatography was performed on an ÄKTA pure™ chromatography system. Analytical reversed-phase HPLC was conducted on an Agilent instrument (1200 series) with a Vydac C18 column (4 × 150 mm, 5 μm), using solvent A (0.1% trifluoroacetic acid (TFA) in water), and solvent B (100% acetonitrile, 0.1% TFA in water) as the mobile phases. Semi-preparative and preparative scale purifications were performed on the Waters 1525 Binary HPLC system using a Vydac C18 semipreparative column (10 mm × 250 mm, 12 μm) at 4 ml/min. ESI–MS analysis was performed on the BioAccord LC-MS system with ESI-Tof based ACQUITY RDa Mass Detector. Cell lysis were performed using a S-450D Branson Digital Sonifier or an Emulsiflex C3. SDS-PAGE and native gel electrophoresis were performed using Bio-Rad electrophoresis systems. Western blot transfer was conducted using the Blot™ 3 Western Blot Transfer Device. SDS PAGE gels, native gels, and Western blots were analyzed with an ImageQuant^Ò^ LAS 7000 S5 imager (Cytiva), Azure 200 Imager (Azure Biosystem), and a LI-COR Odyssey Infrared Imager. Laser microirradiation experiments were conducted using a Marianas and a Zeiss LSM 780. Colony counting was conducted using Oxford Optronix Gelcount^Ò^2. Ionizing radiation was delivered using the CellRadHD X-ray irradiator at 160kV and a dose rate of 2 Gy/min.

#### Software and data analysis

The UCSF Chimera X package was used to obtain structural renderings and models. SnapGene was used to analyze DNA sequences. ImageStudioLite was used for gel densitometry analysis. Kinetic data, gel densitometry data, and immunofluorescence signals were plotted and analyzed using GraphPad Prism (version 10), which was also used to calculate error values and perform t-tests. Grammarly (an AI writing assistant) was used to prepare the manuscript.

#### Expression and purification of recombinant histones and mutants

Recombinant human histones (H2A, H2B, H3.1C96A,C110A, and H4), and histone mutants (H2BS6A, H2BS14A, H2BS6AS14A, H3S10A, H3S28A and H3S10AS28A) were expressed and purified from *E. coli* as previously described with some modifications^1^. Wildtype or mutant expression plasmids (pET30) were transformed into Rosetta(DE3) Competent cells containing the respective gene encoding histones. For protein expression, 5-10 ml of an overnight seed culture was inoculated into 1 L of LB medium supplemented with 100 µg/mL ampicillin or 50 µg/mL kanamycin and grown at 37 °C and 200 rpm until the OD_600_ reached 0.6. Expression was induced with 0.5 mM isopropyl-β-D-thiogalactopyranoside (IPTG), and cultures were incubated for an additional 2–4 h at 37 °C. Cells were harvested and lysed on ice via sonication (2 s on/off cycles for 5 min) in lysis buffer (50 mM Tris-HCl, pH 7.5, 100 mM NaCl, 1mM EDTA, 1mM DTT, 1mM PMSF). The Insoluble lysate was isolated by centrifugation at 28,000g for 20 min, resuspended in lysis buffer, and washed twice under the same conditions. Histone proteins were extracted at room temperature for 1 h using extraction buffer (6 M guanidine hydrochloride, 20 mM Tris-HCl, pH 7.5, 100 mM NaCl, 1 mM EDTA, 1 mM DTT and 1mM PMSF) with gentle end to end nutation. The soluble fraction was obtained after centrifugation at 28,000g rpm for 30 min, and the histone proteins were subsequently purified by preparative RP-HPLC, lyophilized, and stored at -20 °C.

#### Native chemical ligation

Ligation: Hydrazide peptide (H3.1(1-14)K9ac-NHNH2) was synthesized from Fmoc-L-Lys(Boc)-NHN=Pyv Resin (Iris biotech)^2^ following a previously described method with some modifications^3^. Briefly, the acyl hydrazide peptide (2-4 eq) was dissolved in 300 mL oxidation buffer (100 mM K_2_HPO_4_/KH_2_PO_4_, pH 3.0, 6 M guanidine HCl) and kept at -20 °C for 20 min. 40 mL of 1M NaNO_2_ was added to the reaction dropwise in 2-5 min, then let the reaction run for 30 min at -20 °C. Trifluoroethanethiol (TFET, 60-150 eq) was dissolved in cold 300 mL thiol buffer (pH=8.5-9.0, 200 mM K_2_HPO_4_/KH_2_PO_4_, 6 M guanidine HCl) and carefully added to the reaction mixture. After adding TFET to the reaction, monitor the pH using a pH meter and carefully add 1 M NaOH to adjust the pH to approximately 7.5. The reaction was maintained at room temperature for 30 minutes to form the thioester. After 30 min, the reaction was transferred to a tube containing lyophilized truncated histone (1 eq) and sonicated for 2 min. Adjust the pH of the reaction carefully to 7.5 and allow it to proceed overnight. Reaction progress was followed by analytical HPLC and LCMS. After the overnight reaction, the reaction mixture was dialyzed twice in a degassed buffer (100 mM K_2_HPO_4_/KH_2_PO_4_, 6 M guanidine HCl, 0.5 mM Methionine, 0.5 mM TCEP (add fresh); pH approximately 5.5) to remove residual NaNO2.

Desulfurization: Stock solutions of glutathione, Tris(2-carboxyethyl) phosphine (TCEP) and the radical initiator VA44 were added to the ligation mixture for final concentrations of 5 M guanidine HCl, 250 mM TCEP, 100 mM K_2_HPO_4_/KH_2_PO_4_, 20 mM VA44, 40 mM glutathione. The desulfurization reaction was allowed to proceed at 37°C for 16 to 20 hours. Reaction progress was monitored by analytical HPLC and LCMS. Desulfurized histone product was purified by HPLC (20-65% HPLC solvent B gradient over 65 minutes). Fractions were analyzed by LCMS. Pure fractions were pooled, lyophilized, and stored at -20°C.

#### Labeling of histone

H2A was labeled with Cyanine-5 maleimide (Lumiprobe) at position T120C according to previously described protocols^4^.

#### Assembly of histone complexes

Histone octamers, tetramers, and dimers were reconstituted following a previously established protocol^5^ using lyophilized histones. Each histone protein was solubilized in unfolding buffer (6 M guanidine hydrochloride, 20 mM Tris-HCl, pH 7.5, 10 mM DTT) and combined at specific molar ratios: H2A:H2B:H3:H4 = 1.1:1.1:1:1 for octamers, H3:H4 = 1:1 for tetramers, or H2A:H2B = 1:1 for dimers. The mixture underwent stepwise dialysis against refolding buffer (2 M NaCl, 10 mM Tris-HCl, pH 7.5, 1 mM EDTA, 1 mM DTT, 0.2 mM PMSF) at 4 °C with three buffer changes over 24 hours to promote refolding and complex formation. Refolded complexes were centrifuged twice at 5,000 rpm for 10 minutes to remove insoluble materials. The supernatant containing histone complexes was concentrated to 300 µL using Amicon® Ultra filters (30 kDa MWCO) and further purified by size-exclusion chromatography on a Superdex 200 Increase 10/300 GL column equilibrated with refolding buffer. Fractions corresponding to histone octamers, tetramers, or dimers were pooled, concentrated to 2–3 mg/mL, and stored in 50% glycerol at –20 °C for later use.

#### DNA preparation

DNA fragments containing a modified Widom 601 sequence were amplified from a parent plasmid harboring the 147-base pair 601 sequence using Q5 polymerase with 1X Q5 Buffers. The PCR product was precipitated using ethanol precipitation and then purified with a Qiagen miniprep kit. The quality of DNA was assessed by native polyacrylamide gel electrophoresis (5.5% acrylamide gel, 0.5×TBE, 90 V, 65 min) via SYBRÒ gold staining.

Sequence of purified DNAs:

-40N30bio (-5H30bio): gtaatccccttggcggttaaaacgcgggggacagcgcgtacgtgcgtttaagcggtgctagagctgtctacgaccaattgagcggCTGCAGcaccgggattctccagcatcagagacctagggtgatatcagatctg

-35N30bio (0H30bio):

agggagtaatccccttggcggttaaaacgcgggggacagcgcgtacgtgcgtttaagcggtgctagagctgtctacgaccaattgagcggCTGCAGcaccgggattctccagcatcagagacctagggtgatatcagatctg

-30N30bio (5H30bio):

agactagggagtaatccccttggcggttaaaacgcgggggacagcgcgtacgtgcgtttaagcggtgctagagctgtctacgaccaattgagcggCTGCAGcaccgggattctccagcatcagagacctagggtgatatcagatctg

-25N30bio (10H30bio):

cctggagactagggagtaatccccttggcggttaaaacgcgggggacagcgcgtacgtgcgtttaagcggtgctagagctgtctacgaccaattgagcggCTGCAGcaccgggattctccagcatcagagacctagggtgatatcagatctg

-20N30bio (15H30bio):

acgtgcctggagactagggagtaatccccttggcggttaaaacgcgggggacagcgcgtacgtgcgtttaagcggtgctagagctgtctacgaccaattgagcggCTGCAGcaccgggattctccagcatcagagacctagggtgatatcagatctg

-15N30bio (20H30bio):

ctgacacgtgcctggagactagggagtaatccccttggcggttaaaacgcgggggacagcgcgtacgtgcgtttaagcggtgctagagctgtctacgaccaattgagcggCTGCAGcaccgggattctccagcatcagagacctagggtgatatcagatctg

-10N30bio:

tatatctgacacgtgcctggagactagggagtaatccccttggcggttaaaacgcgggggacagcgcgtacgtgcgtttaagcggtgctagagctgtctacgaccaattgagcggCTGCAGcaccgggattctccagcatcagagacctagggtgatatcagatctg

-5N30bio:

atgtatatatctgacacgtgcctggagactagggagtaatccccttggcggttaaaacgcgggggacagcgcgtacgtgcgtttaagcggtgctagagctgtctacgaccaattgagcggCTGCAGcaccgggattctccagcatcagagacctagggtgatatcagatctg

0N30bio:

acaggatgtatatatctgacacgtgcctggagactagggagtaatccccttggcggttaaaacgcgggggacagcgcgtacgtgcgtttaagcggtgctagagctgtctacgaccaattgagcggCTGCAGcaccgggattctccagcatcagagacctagggtgatatcagatctg

5N30bio:

gcgtgacaggatgtatatatctgacacgtgcctggagactagggagtaatccccttggcggttaaaacgcgggggacagcgcgtacgtgcgtttaagcggtgctagagctgtctacgaccaattgagcggCTGCAGcaccgggattctccagcatcagagacctagggtgatatcagatctg

10N30bio:

tcaccgcgtgacaggatgtatatatctgacacgtgcctggagactagggagtaatccccttggcggttaaaacgcgggggacagcgcgtacgtgcgtttaagcggtgctagagctgtctacgaccaattgagcggCTGCAGcaccgggattctccagcatcagagacctagggtgatatcagatctg

15N30bio:

ctgttcaccgcgtgacaggatgtatatatctgacacgtgcctggagactagggagtaatccccttggcggttaaaacgcgggggacagcgcgtacgtgcgtttaagcggtgctagagctgtctacgaccaattgagcggCTGCAGcaccgggattctccagcatcagagacctagggtgatatcagatctg

20N30bio:

atatcgctgttcaccgcgtgacaggatgtatatatctgacacgtgcctggagactagggagtaatccccttggcggttaaaacgcgggggacagcgcgtacgtgcgtttaagcggtgctagagctgtctacgaccaattgagcggCTGCAGcaccgggattctccagcatcagagacctagggtgatatca gatctg

25N30bio:

atccgatatcgctgttcaccgcgtgacaggatgtatatatctgacacgtgcctggagactagggagtaatccccttggcggttaaaacgcgggggacagcgcgtacgtgcgtttaagcggtgctagagctgtctacgaccaattgagcggCTGCAGcaccgggattctccagcatcagagacctagggtg atatcagatctg

30N30bio:

agtggatccgatatcgctgttcaccgcgtgacaggatgtatatatctgacacgtgcctggagactagggagtaatccccttggcggttaaaacgcgggggacagcgcgtacgtgcgtttaagcggtgctagagctgtctacgaccaattgagcggCTGCAGcaccgggattctccagcatcagagaccta gggtgatatcagatctg

Pro-10N30bio:

tcaccgcgtgacaggatgtatatatcCTGCAGgtgcctggagactagggagtaatccccttggcggttaaaacgcgggggacagcgcgtacgtgcgtttaagcggtgctagagctgtctacgaccaattgagcggcctcggcaccgggattctccagcatcagagacctagggtgatatcagatctg

Dis-10N30bio:

tcaccgcgtgacaggatgtatatatctgacacgtgcctggagactagggagtaatccccttggcggttaaaacgcgggggacagcgcgtacgtgcgtttaagcggtgctagagctgtctacgaccaattgagcggCTGCAGcaccgggattctccagcatcagagacctagggtgatatcagatctg

The 601 sequence is underlined.

PstI site (within 601 sequence) is highlighted with capital letters

All sequences (xN30 and xH30) are made with a common biotinylated oligo (/5Biosg/cagatctgatatcaccctaggtctc)

Pro-bio30N10:

agtggatccgatatcgctgttcaccgcgtgacaggatgtatatatctgacacgtgcctggagactagggagtaatccccttggcggttaaaacgcgggggacagcgcgtacgtgcgtttaagcggtgctagagctgtctacgaccaattgagcggCTGCAGcaccgggattctccagcatcagagac

Dis-bio30N10:

agtggatccgatatcgctgttcaccgcgtgacaggatgtatatatcCTGCAGgtgcctggagactagggagtaatccccttggcggttaaaacgcgggggacagcgcgtacgtgcgtttaagcggtgctagagctgtctacgaccaattgagcggcctcggcaccgggattctccagcatcagagac

bio0H10:

agggagtaatccccttggcggttaaaacgcgggggacagcgcgtacgtgcgtttaagcggtgctagagctgtctacgaccaattgagcggCTGCAGcaccgggattctccagcatcagagac

bio15H10:

acgtgcctggagactagggagtaatccccttggcggttaaaacgcgggggacagcgcgtacgtgcgtttaagcggtgctagagctgtctacgaccaattgagcggCTGCAGcaccgggattctccagcatcagagac

bio20H10:

ctgacacgtgcctggagactagggagtaatccccttggcggttaaaacgcgggggacagcgcgtacgtgcgtttaagcggtgctagagctgtctacgaccaattgagcggCTGCAGcaccgggattctccagcatcagagac

The 601 sequence is underlined.

PstI site (within 601 sequence) is highlighted with capital letters

Sequences (xN30bio and xH30bio) are made with a common biotinylated oligo (/5Biosg/cagatctgatatcaccctaggtctc)

Sequences (bio30Nx) are made with a common biotinylated oligo (/5Biosg/agtggatccgatatcgctg) bio0H10, bio15H10 and bio20H10 are made with /5Biosg/agggagtaatccccttggc, /5BiosG/ acgtgcctggagactag, /5Biosg/ctgacacgtgcctgga respectively.

0S0:

TATGCTGCTTGACTTCGGTGATCGGACGAGAACCGGTATATTCAGCATGGTATGGTCGTAGGCTCTTGCTTGATGAAAGTTAAGCTATTTAAAGGGTCAGGGATGTTATGACGTCATCGGCTTATAAATCCCTGGAAGTTATTCGT

0L0:

GGTGAGTATTAACATGGAACTTACTCCAACAATACAGATGCTGAATAAATGTAGTCTAAGTGAAGGAAGAAGGAAAGGTGGGAGCTGCCATCACTCAGAATTGTCCAGCAGGGATTGTGCAAGCTTGTGAATAAAGACACATACTTC

0S30:

CTATGCTGCTTGACTTCGGTGATCGGACGAGAACCGGTATATTCAGCATGGTATGGTCGTAGGCTCTTGCTTGATGAAAGTTAAGCTATTTAAAGGGTCAGGGATGTTATGACGTCATCGGCTTATAAATCCCTGGAAGTTATTCGTTGGAATTCCTCGCGATATCGGATCCACTAG

2L33:

GCATAAGTTAAGTGGTATTAACATATCCTCAGTGGTGAGTATTAACATGGAACTTACTCCAACAATACAGATGCTGAATAAATGTAGTCTAAGTGAAGGAAGAAGGAAAGGTGGGAGCTGCCATCACTCAGAATTGTCCAGCAGGGATTGTGCAAGCTTGTGAATAAAGACACATACTTC AT

#### DNA hybridization

DNA oligo were prepared from ssDNA (standard desalting grade, IDT). The DNA oligo stocks (100 µM) were prepared in 0.1xTE buffer and stored at 4°C until needed. To hybridize, the forward and reverse sequence were combined in stoichiometric amounts at 40 µM concentration in 0.1xTE and heated at 95°C for 5 min and slowly cooled to room temperature (-0.5 °C/min) in a thermocycler. The hybridized adapter DNAs were stored at 4°C.

Oligos for making quenching DNAs are shown below: Fragment FW: /5Phos/acaggatgtatatatctgacacgtgcctggagactagggagtaatccccttggcggttaa Fragment REV: /5Phos/ttaaccgccaaggggattactccctagtctccaggcacgtgtcagatatatacatcctgt

#### Generation of nicked DNA

To generate nicked 30N30 DNA, the DNA fragment were digested by Nt.AlwI (NEB) following the manufacturer’s protocol. The digested DNA was purified via a Qiagen miniprep spin column protocol. Finally, the DNA was ethanol precipitated and resuspended to the desired DNA concentration for nucleosome assembly. The DNA digestion was monitored by native polyacrylamide gel electrophoresis (7 % acrylamide gel, 0.5×TBE, 120 V, 75 min) via SYBRÒ gold staining.

30N30

agtggatacgaGGATCatgt**|**tcaccgcgtgacaggatgtatatatctgacacgtgcctggagactagggagtaatccccttggcggttaaaacgcgggggacagcgcgtacgtgcgtttaagcggtgctagagctgtctacgaccaattgagcggctgcagcaccgggattctccagcatcagagaccta gggtgatatcagatctg

The recognition sequence for Nt.AlwI nicking endonuclease is capitalized (GGATC), and the nicking site is highlighted as **|** symbol.

#### Reconstitution of nucleosomes and hexasomes

nucleosomes and hexasomes were reconstituted using a previously described method^5^.

#### Expression and purification of PARP1

Full-length human PARP1 (114 kDa) was expressed and purified as previously described^6^, with minor modifications. The plasmid containing the human PARP1 gene was transformed into E. coli Rosetta (DE3) competent cells. To prevent auto-modification of PARP1, 1 mM benzamide was added to 2 L of LB medium during inoculation with a 1% (v/v) overnight seed culture. The culture, supplemented with 100 µg/mL ampicillin and 34 µg/mL chloramphenicol, was grown at 37 °C and 200 rpm until reaching an OD₆₀₀ of 0.45, then shifted to 18 °C for 30 min prior to induction with 0.2 mM isopropyl-β-D-thiogalactopyranoside (IPTG). Cells were incubated for an additional 16–24 h at 18 °C, harvested by centrifugation at 10,000 rpm for 10 min at 4 °C, and stored at –20 °C. Frozen cells (2 L) were thawed on ice for 1 h and resuspended in 30 mL of lysis buffer (30 mM HEPES, pH 8.0, 2000 mM NaCl, 20 mM imidazole, 0.1 mM EDTA, 10% glycerol, 3 mM β-mercaptoethanol, 1 mM benzamide, 0.2 mM PMSF, and one protease inhibitor cocktail tablet). Cells were lysed using an Emulsiflex-C3 homogenizer (three passes) and centrifuged at 28,000g for 20 min to remove cell debris. The supernatant was incubated with Ni-NTA resin for 1 h at 4 °C with gentle nutation, followed by extensive washing with 20 column volumes of lysis buffer (three times) and 5 column volumes of wash buffer (30 mM HEPES, pH 8.0, 2000 mM NaCl, 40 mM imidazole, 10% glycerol, 3 mM β-mercaptoethanol, 0.2 mM PMSF, 0.1 mM EDTA). Bound protein was eluted using elution buffer (30 mM HEPES, pH 7.5, 2000 mM NaCl, 0.1 mM EDTA, 300 mM imidazole, 10% glycerol, 1 mM DTT, and 0.2 mM PMSF), and 5 mL fractions were collected. Eluted fractions were analyzed by absorbance at 280 nm, pooled, and treated with HRV 3C protease (25 IU), followed by overnight dialysis at 4 °C in 10 kDa MWCO SnakeSkin dialysis tubing against dialysis buffer (30 mM HEPES, pH 7.5, 150 mM NaCl, 0.1 mM EDTA, 10% glycerol, 1 mM DTT, and 0.2 mM PMSF). Protease cleavage efficiency was verified by 4–20% SDS-PAGE. The cleaved sample was concentrated to 5 mL using Amicon® Ultra centrifugal filters (30 kDa MWCO) and loaded onto a pre-equilibrated 5 mL HiTrap™ Heparin HP affinity column (Cytiva) using an ÄKTA Explorer system at 2 mL/min in Buffer A (30 mM HEPES, pH 7.5, 150 mM NaCl, 0.1 mM EDTA, 10% glycerol, 1 mM DTT). Bound proteins were eluted with a linear gradient from 0–100% Buffer B (30 mM HEPES, pH 7.5, 2000 mM NaCl, 0.1 mM EDTA, 10% glycerol, 1 mM DTT) over 15 column volumes. Eluted fractions were analyzed by SDS-PAGE, and fractions containing PARP1 were pooled and subjected to size-exclusion chromatography using a Superdex 200 Increase 10/300 GL column equilibrated with Buffer A. The purified protein fractions were analyzed again by SDS-PAGE, pooled, and concentrated to ∼20 µM using Amicon® Ultra filters (30 kDa MWCO). Protein concentration was determined by densitometric analysis using a BSA standard curve. To confirm that the purified PARP1 was not auto-PARylated, a Western blot was performed using an anti-PAR antibody, which verified the absence of PAR modification. The final protein was flash-frozen in liquid nitrogen and stored at –80 °C.

#### Expression and Purification of PARP2 and HFP1

PARP2 and HPF1 were expressed and purified as previously described^7^.

#### Expression and Purification of PARG

The catalytically active domain of human PARG (60 kDa) was expressed and purified as previously described with minor modifications^8^. The plasmid encoding PARG was transformed into *E. coli* Rosetta (DE3) competent cells, and 1% (v/v) of an overnight seed culture was inoculated into 1 L of LB broth supplemented with 100 µg/mL ampicillin and 34 µg/mL chloramphenicol. Cultures were grown at 37 °C with shaking at 200 rpm until reaching an OD₆₀₀ of 0.5, then cooled to 16 °C for 30 min before induction with 0.2 mM isopropyl-β-D-thiogalactopyranoside (IPTG). Following 16–24 h incubation at 18 °C, cells were harvested by centrifugation at 10,000 rpm for 10 min at 4 °C and resuspended in 30 mL of lysis buffer (40 mM HEPES, pH 8.0, 300 mM NaCl, 0.1 mM EDTA, 10% glycerol, 3 mM β-mercaptoethanol, 0.2 mM PMSF, and one protease inhibitor tablet). Cells were lysed using an Emulsiflex-C3 homogenizer (three passes) and then clarified by centrifugation at 28,000 × g for 20 min. The supernatant was incubated with 5 mL glutathione agarose resin (2.5 column volumes per liter of culture) for 1.5 h at 4 °C, followed by washing with 20 column volumes of lysis buffer three times. The resin was then equilibrated with digestion buffer (40 mM HEPES, pH 7.5, 150 mM NaCl, 0.1 mM EDTA, 10% glycerol, 1 mM DTT) and incubated with TEV protease (25 IU) in 25 mL digestion buffer for on-bead cleavage overnight at 4 °C on an end-to-end rotator. Cleavage efficiency was assessed by 4–20% SDS-PAGE. The cleaved PARG protein was eluted while the GST tag remained bound to the resin. The eluate was concentrated to ∼300 µL using Amicon® Ultra centrifugal filters (10 kDa MWCO) and further purified by size-exclusion chromatography on a column pre-equilibrated with final buffer (40 mM HEPES, pH 7.5, 200 mM NaCl, 0.1 mM EDTA, 10% glycerol, 1 mM DTT). Eluted fractions were analyzed by SDS-PAGE, pooled, concentrated using Amicon® Ultra filters, quantified by a BSA densitometric assay, flash-frozen in liquid nitrogen, and stored at –80 °C.

#### Electrophoretic mobility shift assay (EMSA)

##### PARP1-mediated chromatin remodeling and modification

Nucleosomes (100 nM), PARP1 (200 nM) with and without HPF1 (10 nM) were incubated at 30 °C in reaction buffer (20 mM HEPES, pH 7.5, 100 mM NaCl, 10% glycerol, TCEP 0.2 mM, and 0.01% (v/v) IGEPAL CA-630). Reactions were initiated with the addition of NAD+ (0.5 mM), and timepoints were quenched with 1x vol of pre-chilled DNA quenching buffer (dsDNA fragment 3 μM, sucrose 10%, 10mM TEK). Time points were kept on ice and resolved by native polyacrylamide gel electrophoresis (PAGE) (5% or 7% polyacrylamide gel in 0.5× TBE). A quenching buffer supplemented with 25 μM olaparib was also tested but showed no difference.

##### PARP1-mediated chromatin disassembly followed by PstI digestion and EMSA of digested chromatin

PARP1-mediated reaction mixture was mixed with an equal volume of restriction enzyme buffer (40 mM Hepes, pH 7.75, 120 mM KCl, 20 mM MgCl2, 25 μM Olaparib, 20% glycerol). PstI 100U/ul was then added to the final concentration of 1.25 U/ul. The reaction mixture was incubated at 30 °C for 30 min before quenching with 2.5 μL of a 20 μM DNA fragment (a final concentration of 2 μM quenching DNA). The reaction mixture was kept on ice and resolved by native PAGE (5 or 7% polyacrylamide gel, 0.5× TBE).

##### PARP1-mediated chromatin disassembly followed by PstI digestion, proteinase K digestion, and native gel analysis of digested DNAs

PARP1-mediated reaction mixture was mixed with an equal volume of restriction enzyme buffer (40 mM Hepes, pH 7.75, 120 mM KCl, 20 mM MgCl2, 25 μM Olaparib, 20% glycerol). PstI 50 U/ul was then added to the final concentration of 4 U/ul. The reaction mixture was incubated at 30 °C for 60 min before quenching with 1.5x volume of proteinase K buffer (10% glycerol, 20 mM HEPES, pH 7.75, 2% SDS, 0.2 mg/ml bromophenol blue, 0.2 mg/ml proteinase K (Invitrogen)). Protein digestion was conducted at 37°C for 1 hour. The resulting digested DNA was resolved by native PAGE (7% polyacrylamide, 0.5× TBE).

##### PARP1-mediated chromatin disassembly followed by PARG-mediated digestion of the PAR chain and EMSA

The PARP1-mediated reaction mixture was quenched with 20 uM olaparib. The reaction was kept on ice for 5 min before adding 0.2x volume of 1.3 uM PARG. The digestion was conducted at 30°C for 30 minutes, followed by quenching with the DNA quenching buffer. The resulting digested DNA was resolved by native PAGE in 7% polyacrylamide, 0.5× TBE.

##### PARP1-mediated chromatin disassembly followed by nucleosome reassembling assay

The PARP1-mediated reaction mixture was quenched with 1x volume of pre-chilled DNA quenching buffer (dsDNA fragment 3 μM, sucrose 10%, 10mM TEK, olaparib 20 μM). The reaction mixture was equilibrated at 30 °C for 10 min before the addition of Cy5-labeled dimer to a final concentration of 0.25 μM. The reaction mixture was incubated at 30 °C for 10 min, then centrifuged at 21,000g for 10 min, and kept on ice. The sample was resolved by native PAGE using 5% or 7% polyacrylamide in 0.5× TBE.

#### Chromatin affinity purification

##### HeLa S3 cell extraction

Nuclear extract was prepared following previously described protocols^9^.

##### Chromatin affinity purification

Biotinylated nucleosome (8.5 pmol at 0.25 uM) in reaction buffer (20 mM HEPES, pH 7.5, 100 mM NaCl, 10% glycerol, DTT 0.5 mM, and 0.01% (v/v) IGEPAL CA-630) was immobilized on beads (Dynabeads Streptavidin Magnetic Beads, Invitrogen; 2 μL bead slurry for 1pmol of nucleosome) by putting on an end-to-end rotor at 4oC for 60 min. PARP1 (1.5 equivalent, 0.375 μM) or a similar volume of buffer was then added to the immobilized beads, and the mixture was further incubated at room temperature on an end-to-end rotor for 30 minutes. NAD+ (0.5 mM) was then added to trigger the disassembly reaction. For the negative control reactions, an equal amount of 1x reaction buffer was added. The reaction mixture was then incubated at room temperature for 15 min on an end-to-end rotator. Quenching DNA was then added to the final concentration of 0.4 μM, and the mixture was incubated at room temperature for 10 min. The supernatant (containing the majority of PARP1 and modified PARP1) was removed, and the beads were washed twice with the reaction buffer. 50ul of NE buffer (20 mM Hepes, 7.9, 100 mM KCl, 0.2 mM EDTA, 0.5 mM DTT, 20% glycerol) was then added to each reaction, followed by the addition of HeLa S3 nuclear extract (16 ul at a concentration of 3.82 ug/ul). The mixture was incubated at 4 °C for 90 minutes on an end-to-end rotor. After removing the supernatant, the beads were washed three times with NE buffer with 10-minute incubation at 4 °C on an end-to-end rotator. Chromatin-bound proteins were eluted with 60 μL of 1x Laemmli SDS buffer. Chromatosome substrates were prepared by mixing biotinylated nucleosomes with H1.0 (5 equivalents) in reaction buffer and incubating for 30 minutes at 37°C before loading onto beads.

#### Western blot protocol

SDS–PAGE gels (4–20% TGX; Bio-Rad) were transferred to nitrocellulose membranes on an iBlot 3 Western Blot Transfer. The membrane was blocked in TBST (50 mM Tris, pH 7.5, 150 mM NaCl, and 0.1% Tween-20) containing 3% milk for 1 h. Membranes were then incubated in primary antibody in PBS containing 3% (w/v) BSA overnight at 4 °C. The membranes were washed three times for 5 minutes with TBST and incubated with the secondary antibody for 1 hour. Membranes were then washed three times for 5 minutes with TBST and then imaged.

#### Mammalian cell culturing and plasmid transfection Cell maintenance protocol

Hela S3 cells were cultured in DMEM (Gibco) supplemented with 10% FBS (Gibco). U2OS cells were cultured in McCoy’s 5A (Modified) Medium (Gibco) supplemented with 10% FBS (Gibco). CHP212 cells were cultured in RPMI media (Gibco) supplemented with 10% FBS (Gibco). Cells were grown in an incubator at 37 °C and 5% CO2. Cells were screened for mycoplasma every 2 months.

#### Plasmid transfection and selection

Plasmid DNA was transfected into U2OS cells using Lipofectamine 2000 (Thermo-Fisher Scientific) according to the manufacturer’s instructions. Cells were selected for 3 weeks with the appropriate selection drug, and single-cell sorting was performed for GFP-expressing cells. pEGFP-N1-PARP1 was a gift from Chris Lord (Addgene plasmid # 211578).

#### Generation of knockout cell line

HPF1−/− U2OS cells were generated using CRISPR technology in the Center for Advanced Genome Engineering at St. Jude. gRNA: CAGE3888.HPF1.9.g6: AAGCGAGUACCCCAAUUCUC PARP1−/− U2OS cells were generated using CRISPR technology in the Center for Advanced Genome Engineering at St. Jude. gRNA: CAGE2961.PARP1.9.g29: GGGACUUUUCCAUCAAACAU Knockout was confirmed by Western blot.

#### Laser microirradiation followed by confocal imaging

##### Laser microirradiation protocol

Laser microirradiation assays were conducted following previously described protocols with some modifications^10,11^. Cells were grown in µ-Slide 8 Well Grid-500 and treated with DMSO or drugs (10 μM) 1 hour before the experiment. Live cells or nuclei were incubated with Hoechst 33342 (Thermo Scientific) at a concentration of 12.3 μg/mL for 15 minutes before measurement. Induced laser microirradiation was performed using laser light at 405 nm on a Marianas SDC microscope (Intelligent Imaging Innovation, Denver CO) using a 63x oil objective. Laser power was set at 4% (ca. 0.29 mW) with photomanipulation settings of 1 msec duration and 1 repetition per ROI. Alternatively, the laser microirradiation assay was performed using a Zeiss LSM 780 confocal microscope (Zeiss, Jena Germany) with a 63x oil objective. Laser power was set at 50% (ca. 1.3 mW) with a 25-50 μsec dwell time. For two-photon laser irradiation using a Chameleon Ti:Sapphire laser, the 750 nm laser power was set to 0.5-0.6% (ca. 4.7 mW) with a 25 μsec dwell time. Both line and square shapes were used to define regions of interest (ROIs). Due to cell movement, square ROIs were primarily used for time-course quantification.

##### Immunofluorescence protocol

At the indicated time point after irradiation, cells were fixed with 4% formaldehyde for 5 minutes at room temperature and washed 3 times with PBS. Cells were permeabilized with PBS containing 0.1% Triton X for 10 minutes at room temperature, then blocked with PBS containing 10% goat serum and RNase A (0.1 ug/ul) for 1 hour at room temperature. Cells were then incubated overnight at 4 °C in the appropriate primary antibody in PBS containing 10% goat serum. Cells were then washed three times for 5 minutes with PBS containing 0.05% Tween 20 and incubated with the secondary antibody for 1 hour. Cells were then washed three times for 5 minutes with PBS containing 0.05% Tween 20 and imaged on a Marianas SDC equipped with a 63x objective lens.

##### m6A labeling

After fixation, permeabilization, and blocking, cells were incubated with 1x CutSmart Buffer (NEB) for 10 minutes at room temperature. The m6A labeling was triggered by exchanging with 1x CutSmart Buffer containing 25 U/ml of EcoGII Methyltransferase (NEB) and 0.32 mM SAM (NEB) at room temperature for 30 minutes. A second dose of SAM (0.32 mM) was then added to the well, and the reaction was continued for an additional 30 minutes. Cells were washed 3 times with PBS containing 0.05% Tween 20 and treated with antibodies according to the above IF protocol. Typically, cells were treated with the m^6^A antibody and imaging before applying the PAR antibodies unless noted in the manuscript.

##### PARG treatment

The poly (ADP-ribose) chain was hydrolyzed by incubation with PARG (0.2 μM) in PBS at room temperature for 1 hour. After PARG treatment, cells can be subjected to m6A labeling or treated with antibodies.

##### Preparation of nuclei for laser microirradiation

Cells were grown overnight in µ-Slide 8 Well Grid-500. The media was removed, and the well was washed with 150 μL of PBS before the addition of 150 μL of 0.5% (w/v) low-melting agarose gel in PBS, which had been pre-equilibrated in a 37 °C heat block for 30 minutes. The mixture plate was left at room temperature for 5 minutes, then transferred to a cold room for 30 minutes to solidify the gel. Cells were permeabilized by incubating twice with 400 μL of PBS containing 0.05% Triton X-100 and protease inhibitor at 4°C for 20 minutes each. Cells were then washed twice with PBS containing protease inhibitor at 4°C every 30 minutes. NAD^+^ was added 5 minutes before conducting laser microirradiation.

##### Dimer-complex formation

dsDNA was incubated at 30 °C in complex-forming buffer (10 mM Tris-HCl, pH 7.5, 1 mM EDTA, 10 mM KCl, 350 mM NaCl, 1 mM DTT, 0.2 mM PMSF, 0.03% (v/v) Triton X). Pre-form histone dimer (1 equivalent, 10 mM Tris-HCl, pH 7.5, 1 mM EDTA,1 M NaCl) was added and mixed quickly make a final 500 mM NaCl. The mixture was incubated at room temperature for 10 minutes. Then, 10TEK (10 mM Tris-HCl, pH 7.5, 1 mM EDTA, 10 mM KCl, 1 mM DTT, 0.2 mM PMSF, 0.03% (v/v) Triton X) was added to a final concentration of 1 μM dimer complex in 100 mM NaCl. The complex can be kept at 4 °C for at least 1 week.

##### In situ reassembly of chromatin

At the indicated time point after irradiation, cells were fixed with 1% formaldehyde for 5 minutes at room temperature and washed 3 times with PBS. Cells were permeabilized in PBS containing 0.1% Triton X for 10 minutes at room temperature, then treated with RNAse (0.1 ug/ul) in blocking buffer (PBS containing 10% fish gelatin) for 1 hour at room temperature. 200 μL of dimer complex (25 nM) in PBS containing 10% fish gelatin was added to each well and incubated at room temperature for 1 hour. Cells were then washed 3 times with PBS containing 0.05% Tween 20 before imaging or antibody treatment.

##### Data processing and analysi

Images were processed using Fiji. The ROIs were manually drawn and used to measure the fluorescence intensity within each stripe in the input (raw) image. Local background was defined as a similar shape within the same cells with the lowest signals. The median of the signal of each ROI was normalized over the median of the corresponding background and plotted using GraphPad Prism (version 10). Statistical comparisons between two conditions were performed using two-tailed Welch’s t-tests in GraphPad Prism (version 10). For the two-photon experiment, time-course imaging was performed, and GFP signals from ROIs were directly acquired on the Zeiss LSM 780. Data were normalized, fitted to a one-phase exponential decay model, and plotted using GraphPad Prism (version 10).

#### Colony-forming assay

Colony-forming assays were performed following previously published methods^12^. Briefly, U2OS cells were seeded into 6-well plates at the specified cell densities indicated in the manuscript using conditioned medium (50% fresh complete medium and 50% previously used medium). Cells were allowed to adhere for 6 hours prior to drug treatment. Cells were then treated with the indicated drugs for 30 minutes. Following treatment, the drug-containing medium was removed, and the cells were washed once with PBS before being replaced with fresh conditioned medium. Cells were cultured for 10 days to allow colony formation, after which colonies were fixed and stained with crystal violet (0.1% v/v).

##### Data processing and analysis

Colony counting was performed using the Oxford Optronix GelCount® 2 system. Colony numbers were divided by the number of cells initially seeded to calculate the percentage survival. Survival percentages were then normalized to the mean of the control condition to obtain normalized survival values (normalized counts). The normalized data were plotted using GraphPad Prism (version 10). Statistical significance was determined using two-way ANOVA followed by Sidak’s multiple comparisons test.

#### Antibodies list

| antibody name | Species | catalog no |
| --- | --- | --- |
| Strep-Tag II Monoclonal Antibody (Mix-mA), Biotin, Bioss | mouse | 50-227-9974 |
| Poly/Mono-ADP Ribose (D9P7Z) Rabbit mAb | rabbit | 89190S |
| Poly(ADP-ribose), mAb (10H) | mouse | ALX804220R100 |
| Phospho-KAP-1 (Ser824) Polyclonal Antibody | rabbit | A300-767A |
| Phospho-Histone H2A.X (Ser139) #2577 | rabbit | 2577 |
| Phospho-Histone H2A.X (Ser139) | mouse | MilliporeSigma™ 05636 |
| PARP-1 N-terminal antibody (pAb) | rabbit | 39061 |
| PARP-1 (F-2) | mouse | sc-8007 |
| N6-Methyladenosine (m6A) Recombinant Rabbit Monoclonal Antibody | rabbit | MA5-35350 |
| Mre11 Antibody (12D7) - Azide and BSA Free | mouse | NB100-473 |
| KU-86/XRCC5 | mouse | sc-5280 |
| KU-70/XRCC6 | mouse | sc-17789 |
| HPF1 (E8B3W) Rabbit mAb | rabbit | #47590 |
| Histone H4 antibody [mAbcam 31830] - ChIP Grade | mouse | ab31830-100ug |
| Histone H3 (1B1B2) Mouse mAb | mouse | 14269S |
| Histone H1.0 antibody [EPR6536] | rabbit | ab134914-100uL |
| HA-Tag (C29F4) Rabbit Monoclonal Antibody #3724 | rabbit | #3724 |
| GAPDH | mouse | sc-365062 |
| DNA-PKcs (G-4) | mouse | sc-5282 |
| DNA Topoisomerase II alpha | mouse | sc-365916 |
| CHD4 (D4B7) Rabbit Monoclonal Antibody | rabbit | D4B7 |
| Anti-Histone H2B antibody | rabbit | ab1790 |

